# PRosettaC: Rosetta based modeling of PROTAC mediated ternary complexes

**DOI:** 10.1101/2020.05.27.119354

**Authors:** Daniel Zaidman, Nir London

## Abstract

Proteolysis-targeting chimeras (PROTACs), which induce degradation by recruitment of an E3 ligase to a target protein, are gaining much interest as a new pharmacological modality. However, designing PROTACs is challenging. Formation of a ternary complex between the protein target, the PROTAC and the recruited E3 ligase is considered paramount for successful degradation. A structural model of this ternary complex could in principle inform rational PROTAC design. Unfortunately, only a handful of structures are available for such complexes, necessitating tools for their modeling. We developed a combined protocol that alternates between sampling of the protein-protein interaction space and the PROTAC molecule conformational space. Application of this protocol - PRosettaC - to a benchmark of known PROTAC ternary complexes results in near-native predictions, with often atomic accuracy prediction of the protein chains, as well as the PROTAC binding moieties. It allowed the modeling of a CRBN/BTK complex that recapitulated experimental results for a series of PROTACs. PRosettaC generated models may be used to design PROTACs for new targets, as well as improve PROTACs for existing targets, potentially cutting down time and synthesis efforts.

## Introduction

Proteolysis targeting chimeras (PROTACs) are modular small molecules that induce the degradation of a target protein^1–3^. PROTACs can be conceptually divided into three parts: A target binding moiety, an E3 ubiquitin-ligase binding moiety and a linker that connects the two. Through simultaneous binding of their target and E3 ligase they induce the formation of a ternary complex that facilitates the ubiquitination of the target which then leads to its proteasomal degradation.

PROTACs offer several advantages over traditional small molecule inhibitors, including: complete inhibition of function, not limited to enzymatic function, or a specific site on the target, long duration-of-action that is proportional to the protein turn-over rate, enhanced selectivity^4–7^ and the ability to work at sub-stoichiometric concentration^8^, since a single PROTAC molecule can degrade multiple copies of the target. These advantages propelled wide interest in their development as chemical tools and potential drugs.

Most reported PROTACs to date are based on a known and potent small-molecule target binder. Typically a binder for which a co-crystal structure was available to define a suitable exit vector for the installation of the linker. From the E3 binding side, the vast majority of PROTACs are limited to just two ligases: CRBN which is targeted by various immunomodulatory drugs (IMiDs) such as thalidomide^9^ , pomalidomide^10^ or lenalidomide^11,12^ and VHL for which high-affinity small molecule ligands were developed ^13,14^.

A key challenge in PROTAC design to date is the selection of the optimal linker to connect these two binding components. In most cases, linkers of various lengths are screened using synthetically accessible chemistry^15–19^. In some cases, it was shown that a protein-protein interface between the target and the E3 ligase, including interactions with the linker, is important for cooperativity ^4^. In other cases, it was sufficient for the linker to cross a certain length, and beyond that, longer linkers were also successful^20^. Evidence is accumulating that a stable ternary complex involving protein-protein interactions is important for efficient degradation ^4,21,22^. Still, there is currently no way to predict or design successful linkers, which necessitates significant synthetic effort.

Despite the increasing popularity of PROTACs and the many examples of experimentally validated, cell-active PROTACs, structural information on Target/E3/PROTAC ternary complexes is still very limited. The ability to model such ternary complexes can in theory inform the design of more potent PROTACs, significantly reduce the number of linker designs required to achieve efficient degradation, as well as rationalize the structure activity relationship of already known series of PROTACs. However such modeling efforts are still rare.

A pioneering work in modeling PROTAC mediated ternary complexes^23^ explored four methods to generate ensembles of ternary complexes and had some success in recapitulating the very few available crystal structures of such complexes. Some system specific modeling efforts were also reported. For instance, docking followed by short molecular dynamics simulations were used to model p38 isoforms in complex with PROTACs and VHL and to predict residues that contribute to linker interactions and selectivity of the PROTACs^5,6^. PROTAC ensembles in the absence of the protein complex were used to define distance distribution between the binding partners^24^. HADDOCK^25^ was used to dock Sirt2 and CRBN^26^, subsequent docking of a PROTAC into the complex recapitulated the known binding modes of the separate protein binding moieties. Notably, RosettaDock was used by Nowak et al. ^27^ to inform the design of selective PROTACs. Two relevant computational approaches for linker design were recently introduced ^28,29^ that can be used to design PROTAC linkers based on the protein-protein binary complex. While these may be suitable as a sub-procedure within a general protocol, they cannot in and of themselves model ternary complexes.

Here we present PRosettaC, a holistic protocol for modeling PROTAC mediated ternary complexes. It combines global docking with PatchDock^30,31^ under PROTAC derived distance constraints, and local docking with RosettaDock^32^, followed by modeling of the PROTAC into the ternary complex ^33^. The protocol was able to accurately recover published structures of ternary complexes to near atomic resolution, and recapitulate experimental trends for a couple of model systems. This general protocol should be useful in the design of new PROTACs for a wide variety of targets.

## Results

### The PRosettaC protocol

The protocol includes several consecutive steps, each intended to decrease the large conformational search space which includes the protein-protein interaction degrees of freedom, the PROTAC internal conformation and its pose relative to the protein-protein complex (Fig. 1).

**Figure 1.**
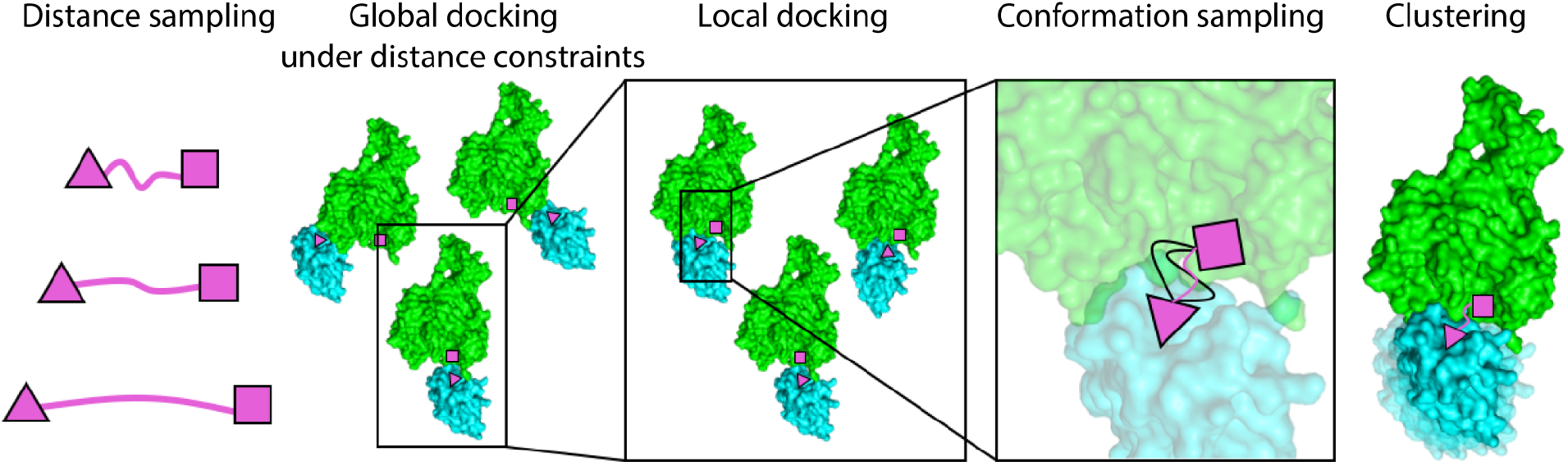
An overview of the PRosettaC protocol. The protocol consists of the following consecutive steps: **1.** Sampling of the distance between the two ligand anchor points. **2.** Constrained global protein-protein docking with PatchDock. **3.** Local docking with RosettaDock. **4.** Generating constrained PROTAC conformations compatible with the local docking solutions. **5.** Clustering of the top scoring results.

The input for the protocol are structures, or models, of the protein target and E3 ligase, each in complex with its own binder, as well as a smiles string, representing the entire PROTAC (Fig. 2). Similar to Drummond and Williams^23^, we use two anchor atoms in the two binders which are defined as part of the input. In most cases we chose the anchor atoms as the atoms through which the PROTAC’s linker is attached to the two binders, with the exception of thalidomide, where the attachment atom is not uniquely defined within the SMILES string, and thus we chose a more central atom.

**Figure 2.**
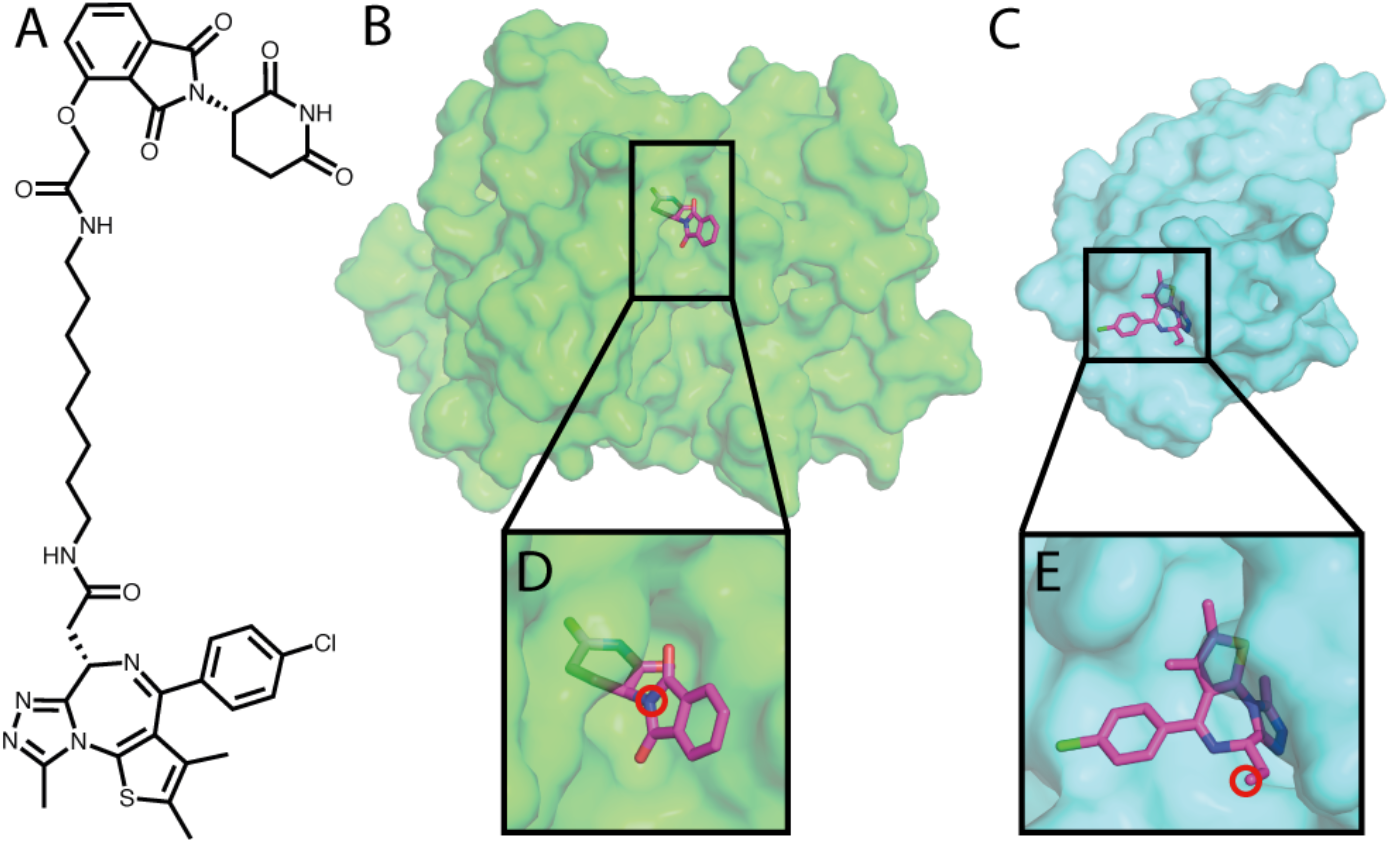
The input for the PRosettaC protocol. **A.** The chemical structure of a BRD4 PROTAC molecule including a CRBN binder (thalidomide), a BRD4 binder (JQ1) and an alkane linker. **B.** Structure of CRBN with thalidomide. **C.** Structure of BRD4 with JQ1. **D.** Thalidomide with the anchor atom marked. **E.** JQ1 with the anchor atom marked. All structures are based on PDB: 6BOY.

#### Step 1. Rough sampling of the distance between the anchor points

To estimate the distance between the anchor atoms we do the following: for each value starting from 1Å and in increments of 1Å we generate 200 random pairs of binder positions with the anchor distance set to that value. Then, for each pair of binders positions, we generate a random conformation of the entire PROTAC which incorporates the ‘fixed’ binder positions. Due to geometrical constraints of the molecule, some of the pairs fail to generate a valid PROTAC conformation while others succeed. We sum up the successfully generated PROTAC conformations for each distance bin (Fig. 3A) , and then pick distance constraints according to the distribution of successful conformations (see methods, Supp. Fig. 1 and Supp. Table 1 for detailed explanation of the choice of distance constraints).

**Figure 3.**
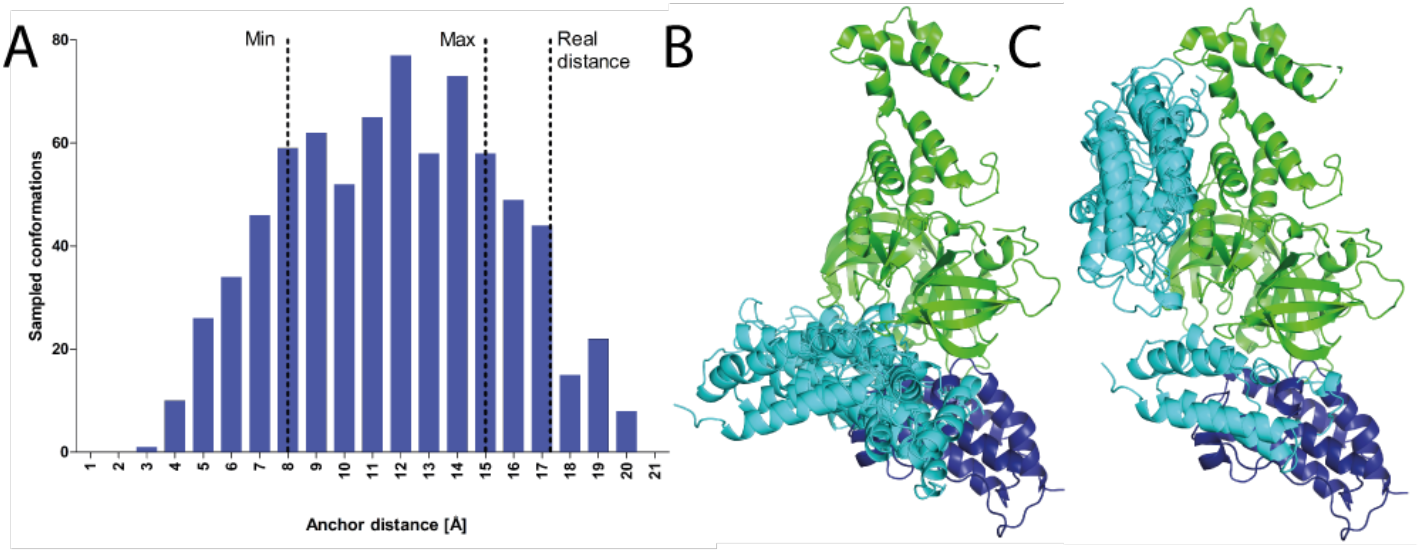
Sampling the PROTAC anchor points distance distribution successfully constrains global docking. **A.** For each distance (in 1Å increments), we generate 200 random orientations of the target and E3 binders, and try to generate PROTAC conformations to match these random orientations. The histogram represents successfully generated conformations of an example PROTAC (PDB: 6BOY). Based on this histogram the constraints chosen for the following step were 8-15Å. Despite the crystallographic distance falling outside the constraints boundaries in this example, the protocol eventually is able to recapitulate the correct binding mode. See Supp. Fig. 1 and Supp. Table 1 for more details on boundary selection, and the crystallographic distances. **B.** Top 5 solutions of global protein-protein docking, constrained by the range calculated by the previous step of the protocol. **C.** Top 5 solutions of unconstrained protein-protein docking. Green: E3 ligase CRBN, dark blue: crystallographic BRD4 (6BOY), cyan: docked BRD4.

#### Step 2. Global protein-protein docking

We use PatchDock, a very efficient global docking algorithm ^30^, to sample the protein-protein interaction space. The distance between the two anchors is forced to be within the constraints calculated in the previous step. This limits the docking search space considerably (Fig. 3B,C).

#### Step 3: Local docking refinement

After using PatchDock to rapidly sample the protein-protein interaction space, we use RosettaDock ^34^ local docking to produce 50 high-resolution models for each PatchDock global docking solution.

#### Step 4: Modeling the PROTACs into the docking solutions

We treat each of the local docking solutions as a hypothesis for the final ternary complex (Fig. 4A). Thus, we constrain the binders to be in the appropriate positions, derived from the protein-protein docking (Fig. 4B), and construct 100 conformations of the full PROTAC that would fit these positions (Fig. 4C). In other words, we create up to 100 random linker conformations which could connect the two binders, given their position in the docking solution. For a significant portion of the local docking solutions, after 100 trials, no conformations could be found. Thus, no linker can attach the two binders in these conformations (Supp. Table 1). For example, for PDB: 6BOY, out of ~36,000 local docking solutions, only ~28,000 got at least one generated conformation (Supp. Table 1).

**Figure 4:**
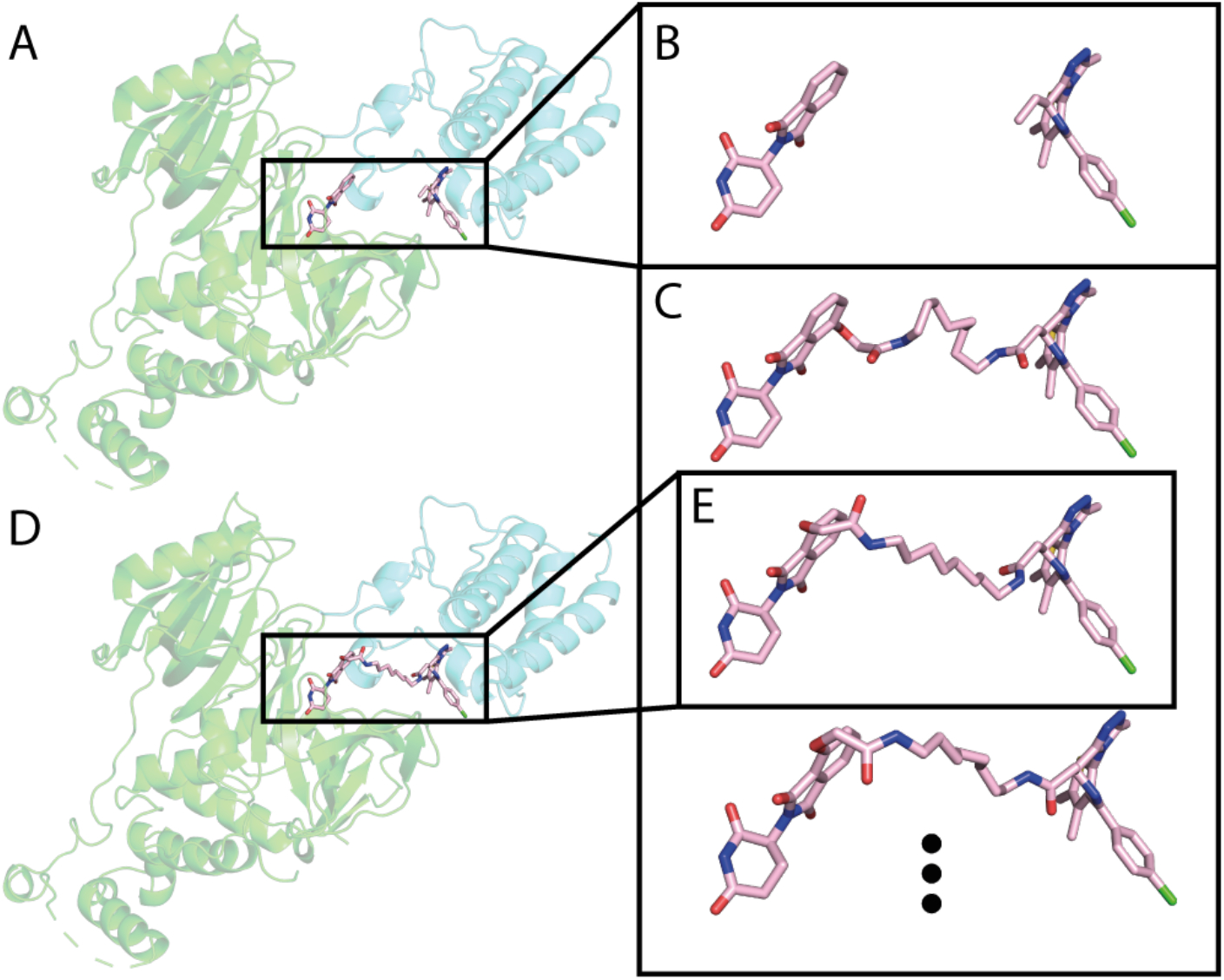
Generating PROTAC conformations in the context of the protein-protein interaction. Using RDKit, we generate up to 100 PROTAC conformations, to match the binders orientation of each local docking solution. Then, we use Rosetta Packer to choose the best conformation to fit the protein-protein docking model. **A.** An example local docking solution for PDB: 6BOY. **B.** The binders orientation, extracted from the local docking solution. **C.** Constrained conformations that bridge between the binders orientation. **D.** Output model of Rosetta after choosing the best constrained conformation. **E.** The conformation chosen by Rosetta. Green: E3 ligase CRBN, cyan: predicted BRD4, light pink: predicted PROTAC conformation.

We use the Rosetta Packer ^35^, to choose the optimal linker conformation of those generated (if one exists) that could connect the two binders in the context of the docking solution (Fig. 4E). This step outputs possible ternary complexes (Fig. 4D) and prevents clashes between the PROTAC and the protein-protein complex. At this point we filter complexes with high Rosetta energy score, and exclude them from further analysis.

#### Step 5: Clustering top scoring complexes

After scoring the ternary complexes from the last step, we cluster them, under the assumption that near-native solutions will be sampled many times. From the 1,000 models with the lowest Rosetta score, we choose the 200 with the best interface score (energy of the complex less the energy of the separate components after side-chain minimization). We cluster these 200 complexes with a threshold of 4Å RMSD for the moving chain using the DBSCAN ^36^ clustering method, and rank the clusters by the number of models in each cluster. Between clusters of the same size, the ranking is based on the average score of the final models. The 4Å RMSD threshold was chosen based on manual inspection of similarity with different thresholds (Supp. Fig 2), and is consistent with accepted criteria in protein-protein docking^37^. The “near-native’ cluster is defined as the top cluster which contains at least one near-native structure - a structure with Cα RMSD below 4Å to the crystallographic conformation.

### Optimization of hyper parameters

The protocol contains several parameters that require optimization. These include: choosing whether to use the entire E3 ligase complex from the crystal structure or only the CRBN or VHL monomer; the RMSD threshold for clustering the global docking results; the number of top scoring models to consider for clustering; the number of top interface scores within the top scoring models. Choosing the optimal parameters was not straightforward since there are only a handful of ternary structures available. We chose to optimize them on a set of five structures which were previously used for benchmarking^23^ (PDB IDs: 6BOY, 6BN7, 6BN8, 6BN9 ^27^, 5T35 ^4^). We tested a matrix of different combinations for the five aforementioned parameters (Supp. Table 2). The choice of docking the E3 monomer or complex had little effect, likely since the distance constraints to the global docking already restrict the possible interaction interfaces, we hence chose to use the monomer to reduce runtime. Increasing the number of RosettaDock local docking models from 5 to 10 to 50 improved performance.

### Accurate recapitulation of ternary complexes using bound structures

With the best combination of parameters we were able to predict near-native models for four out of the five cases in the train set. In all four cases the near-native models were found in the first or second ranked clusters (Table 1; See Fig. 5 for the lowest RMSD solution for each of the four structures).

**Table 1.** PRosettaC performance against known ternary complex structures.

|  | Train |  |  |  |  | Test |  |  |  |  |
| --- | --- | --- | --- | --- | --- | --- | --- | --- | --- | --- |
| PDB | 5T35 | 6BOY | 6BN7 | 6BN8 | 6BN9 | 6BNB | 6SIS | 6HAX | 6HAY | 6HR2 |
| Rank<sup>a</sup> | 2/46 | 2/22 | 1/29 | NA/51<sup>c</sup> | 2/40 | 1/20 | 3/17 | 71/76<sup>d</sup> | 20/35<sup>d</sup> | 18/61 |
| RMSD <10Å<sup>b</sup> | 3.5% | 17.0% | 62.0% | 0.5% | 22.0% | 47.0% | 19.0% | 2.5% | 1.0% | 5% |

<sup>a</sup> Rank of the near-native cluster / total number of clusters.
<sup>b</sup> Percent of models that show less than 10Å RMSD to the crystal structure (performance measure reported by <sup>23</sup>)
<sup>c</sup> No near-native clusters were found for this structure.
<sup>d</sup> These clusters were singletons, i.e. contained a single structure.

**Figure 5.**
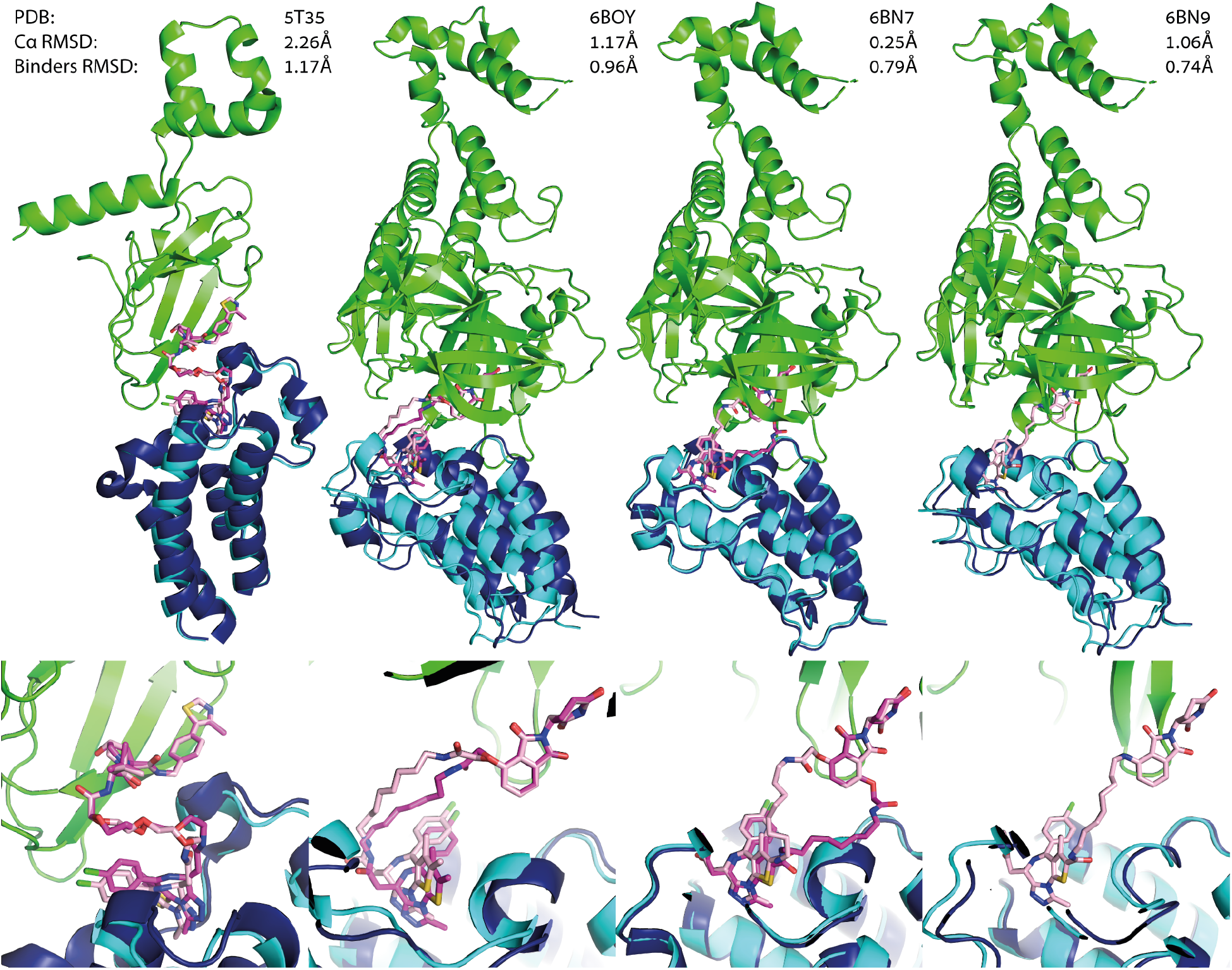
PRosettaC can recapitulate known ternary complexes to sub-angstrom accuracy. Upper panels: near native predictions for four structures of the training set: 5T35, 6BOY, 6BN7 and 6BN9. With reported Cα RMSD of the moving chain (BRD4) and RMSD over the small molecule binders (excluding the linker). Lower panels: zoom in on the ligand binding predictions. In three out of the four, PRosettaC achieved sub-angstrom predictions of the PROTAC “heads’. Green: E3 ligase CRBN, dark blue: x-ray BRD4, cyan: docked BRD4, magenta: x-ray PROTAC conformation, light pink: predicted PROTAC conformation. Note that the PROTAC molecule is not resolved in 6BN9 and was modelled by PRosettaC. The RMSD values for the binders are based on alignment to PDB: 6BN7.

### PRosettaC compares favourably to previously reported methods

Drummond and Williams^23^ reported the results of their methods against five structures. To be able to compare our results, we first had to realign the models, since they chose the degradation target as the static chain, while we chose the E3 ligase, due to the large size of CRBN compared to BRD4. The results (Supp. Table 3) are therefore based on the E3 Cα RMSD. As a metric for success they report the percent of models with RMSD to the x-ray structure below 10Å. In three cases (6BOY, 6BN7 and 6BN9), PRosettaC outperformed their method. In one case (5T35) their method did better. Both methods performed poorly on 6BN8. We should note that while Drummond and Williams were not able to generate successful models for 6BOY, our method generated 17% such models, ranking the native cluster in second place. Also, even though we could not match the overall fraction of near native solutions for 5T35, we did rank the cluster which includes the native structure in second place. To our understanding, Drummond and Williams do not report ranking.

### Application to a new set of test structures

After optimizing the hyper parameters on the training set (Supp. Table 2), we tested PRosettaC on five additional structures which were not modeled in previous work (PDBs: 6BNB, 6SIS, 6HAX, 6HAY, 6HR2), nor optimized against. The most successful out of these was 6BNB, a complex of BRD4 with CRBN (Fig 6A), in which CRBN shows an “open’ conformation, substantially different from the closed conformation seen in most structures. The top ranking cluster, which included 67.7% of the final models, was the near-native cluster. The PROTAC molecule (dBET57) is not modeled in the original structure, likely due to the low resolution of the structure (6.3Å). We placed the binders by aligning structures with the individual binders onto 6BNB (using domains from PDB: 6BOY as templates). Based on our prediction, we were able to easily model the PROTAC (Fig. 6A). A second successful prediction was for a complex of VHL with BRD4 (PDB: 6SIS), bridged by a non-conventional macrocyclic PROTAC^38^. The near native cluster was ranked third, and the ligand binding moieties were recapitulated with sub-angstrom accuracy (Fig. 6B). For the rest of the test cases, all complexes of VHL with SMARCA2/4^39^, while near-native clusters were found, they were not ranked highly (Table 1).

**Figure 6.**
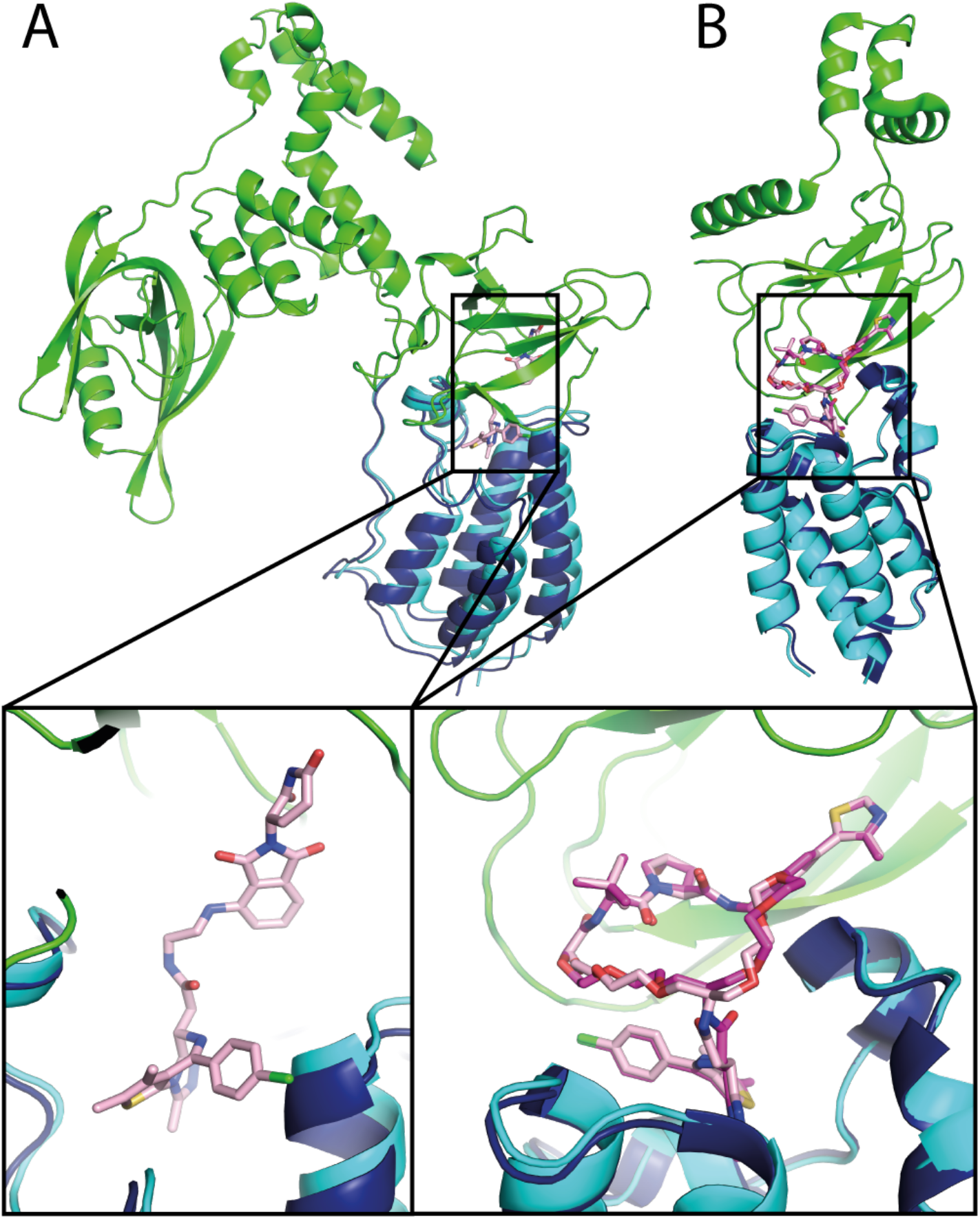
Near native predictions for two non-traditional PROTAC ternary complexes. **A.** Near native prediction of PDB: 6BNB of a BRD4 PROTAC with an open conformation of CRBN. The Cα RMSD is 1.21Å. The native cluster was ranked 1, containing 67.7% of the final models. The PROTAC molecule is not modeled in the original structure, but could be easily modeled in using PRosettaC **B.** Near native prediction of PDB: 6SIS a complex of BRD4 and CRBN with a macrocyclic PROTAC. The Cα RMSD is 1.3Å, with binders RMSD of 0.66Å. The native cluster was ranked 3. Green: E3 ligase; Dark blue: crystallographic BRD4; Cyan: predicted BRD4; Light pink: predicted PROTAC conformation.

### PRosettaC models can help to explain structure activity relationships

The BRD4 triple mutant K378Q, E383V, A384K was experimentally shown to reduce ternary complex formation with VHL^4^ and was previously used as a negative test case^23^. We introduced these three mutations in-silico (based on PDB: 5T35), and re-ran the protocol. While for the wild-type BRD4, the native cluster was ranked second out of 46, containing 9% of the final models, none of 34 clusters proposed for the triple mutant were near-native. This suggests that indeed these three positions were crucial for the formation of the productive ternary complex.

To test our method’s ability to predict unknown ternary complexes, we next turned to a systematic study of PROTACs against BTK ^20^. The authors tested 11 different PROTACs with increasing linker lengths, between 5-21 atoms, for their ability to degrade BTK, as well as for their ability to form a ternary complex (using a TR-FRET experiment). PROTACs 1-4 cannot degrade BTK efficiently, nor lead to ternary complex formation at low concentrations. PROTACs 6-11 were all efficient in both degrading BTK and forming the ternary complex. The authors used computational modeling to recapitulate their experimental results. Using a proprietary pipeline to create models of the ternary complex they were able to produce non-clashing models for PROTACs with longer linkers (PROTACs 7-11), but not for PROTACs with shorter linkers (1-4).

We applied PRosettaC to this model system. We used a structure of CRBN bound to thalidomide (PDB: 4CI1), and BTK bound to the inhibitor on which the PROTACs were based (PDB: 6MNY). Our results (Supp. Table 4) correlated very well with the experimental data. The first distinction between the active and non-active PROTACs was the number of generated models. For PROTACs 1-3 our protocol was not able to generate any ternary structure. For PROTACs 4 and 5, it generated less than 100 models, and for 6-11 more than 2000 models. Second, we noticed that the top ranked clusters of PROTACs 7-9 are very similar, and are also similar to the second ranked cluster of PROTACs 10 and 11. This suggests that the complex represented by these clusters is a low energy solution that may lead to efficient degradation. We pooled together and clustered all of the top final models from the separate runs. Indeed, we found that the most populated cluster is the aforementioned one, which includes representatives from PROTACs 6-11 but not 4 and 5. The second cluster includes representatives of PROTACs 9-11 only, and the third cluster PROTACs 7-11. To further fine-tune our prediction, we clustered this top overall cluster with a lower RMSD threshold of 1Å, resulting in a tight ensemble of 60 similar structures, still containing representatives of PROTACs 6-11 (Fig. 7A).

**Figure 7:**
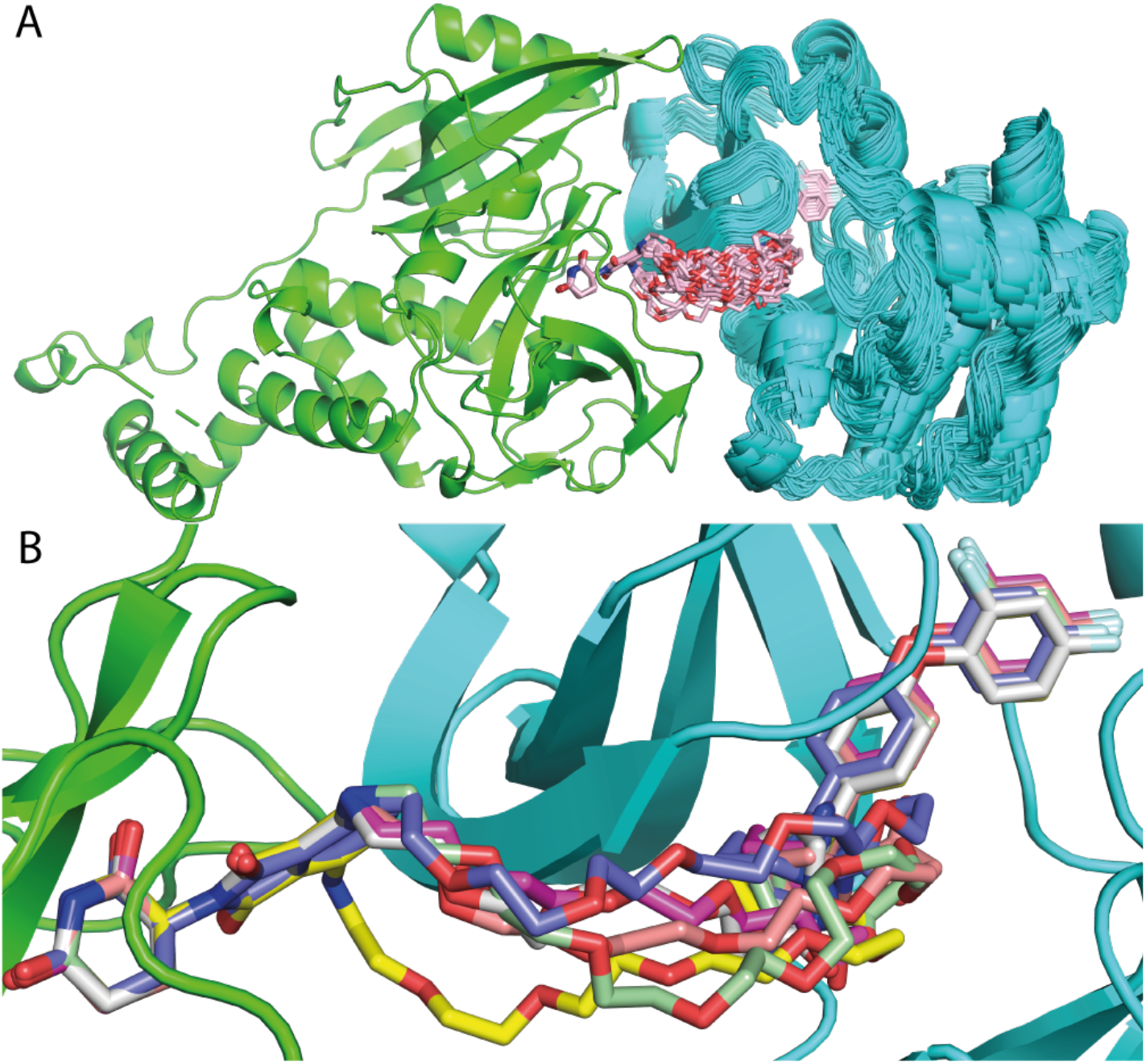
High confidence prediction of a BTK-CRBN ternary complex. Modeling of a BTK-CRBN ternary complex, with a series of 11 PROTACs recapitulates which PROTACs are active and which are not and suggests a high-confidence model for the interaction. **A.** A cluster (with 1Å threshold) of 60 models from various PRosettaC runs of BTK (cyan) in complex with CRBN (green) contains representatives from active PROTACs 6-11. **B.** Representative of PROTACs 6-11: 6 - gray, 7 - magenta, 8 - yellow, 9 - orange, 10 - purple, 11 - light green.

## Discussion

The field of PROTAC mediated targeted degradation is gaining tremendous momentum, as highlighted by the recent positive results from the first PROTAC (ARV-110) to enter clinical trials^40^. Structure-based design is a key paradigm in drug discovery, and PROTAC design can clearly benefit from structural insights, as evidenced by the few cases in which ternary complex structures were actually determined. Gadd et al. used the structure of the BRD4/VHL/MZ1 ternary complex to design a new PROTAC (AT1) in which the linker uses a different exit vector from the VHL ligand^4^. Farnaby et al. used the complex of SMARCA2/VHL/PROTAC-1 complex to rigidify the PROTAC linker and introduce a new pi-stacking interaction, resulting in a significantly improved molecule. In the absence of ternary complex experimental structures, PROTAC designers can turn to modeling in order to generate design hypotheses.

Modeling of PROTAC mediated ternary complexes, however, is a challenging task for multiple reasons. First, is the very small number of available structures of such interactions, which complicates the benchmarking of new methods. Second, is the fact that the E3 ligase and protein target are not cognate interaction partners, thus any binding interface that may be stabilized by the PROTAC is likely transient with weak affinity^41–43^ such low affinity PPIs are known to be challenging for docking methods^44^. Another difficulty is the need to simultaneously model protein-protein interactions and protein-ligand interactions, the latter often with a significant number of rotatable bonds.

Here, we presented a general approach which overcomes some of the challenges by using the PROTAC’s distance distribution to restrict the protein-protein docking search space, and then uses the docking solution to restrict the ligand conformational search space. PRosettaC was very successful in recapitulating the ternary complexes in 6 out of 10 cases. Moreover, the near-native models were ranked in the top three solutions. It was further able to recapitulate trends for a series of BTK targeting PROTACs and propose a high-confidence model for the BTK/CRBN ternary complex, by combining the experimental results from various PROTACs ^20^.

Based on these successful results we suspect that PRosettaC would be a useful tool to enable structure-based PROTAC design. The fact that in most cases the near-native complex was ranked in the top three clusters, suggests that in the future a small number of PROTACs that are designed based on a limited number of the top clusters could be used to validate which model captures the productive complex that leads to degradation.

One design prediction already emerged based on the modeling of the BRD4/CRBN/dBET23 complex (PDB: 6BN7), for which PRosettaC identified the native cluster as the top ranking solution, including 65.5% of the final structures. In this case, the protein-protein interaction was accurately recapitulated, as were the thalidomide and BRD4 binder moieties. However, a flip of the thalidomide phthalimide which scored better, forced the linker to exit out of the opposite atom compared to the one observed in the crystal structure (Fig. 5). Testa et al. previously used the structure of BRD4/VHL/MZ1 elegantly to create a macrocyclic PROTAC^38^.

The 6BN7 modeling result suggests a similar strategy would work for the BRD4/CRBN complex as well (Supp. Fig. 3).

Despite its success, our protocol still suffers from some challenges. In the set of the original five test structures, PRosettaC was least successful for PDB: 6BN8, a complex of BRD4/CRBN and the PROTAC dBET55 for which the native solution did not belong to any of the clusters. dBET55 contains a very long PEG9 linker. The length and flexibility of the linker presents a problem for the anchor distance sampling step, and later to global docking, which runs with very loose constraints. Indeed, global docking produced an exceptionally large number of structures, and most near-native solutions were not ranked among the top 1,000 (Supp. Fig 4). Since such long and flexible linkers are rare for PROTACs, we don’t consider this a significant failure. For another three cases involving complexes of VHL with SMARCA2/4, PRosettaC was able to sample near-native solutions but did not rank them as one of the top clusters. Analysis of the global docking results (Supp. Fig. 4) suggests that not sampling a near-native conformation for these complexes in the global docking step, may be the reason for these failures. Future developments may investigate optimization or alternatives to the global docking step. One possibility is that the crystal structure represents a single possible conformation for the interaction, stabilized by crystal contacts (Supp. Fig. 5), while other conformations proposed by our protocol may be sampled in the solution state. Indeed, Nowak et al. used such a “non-native’ docking prediction to guide the design of a new and more selective PROTAC^27^, strongly suggesting that such additional conformations are sampled in solution and can be accessed by a suitable PROTAC. Hence, even if the top predicted clusters do not correspond to the crystal structure, they may still represent attractive opportunities for PROTAC design.

Another important caveat of this (and previous^23^) benchmark is the use of the bound structures. Ideally the protocol should be able to reproduce the ternary complex starting from the structures of the unbound monomers. Currently PRosettaC fails to rank near-native clusters when starting from unbound structures. Future improvements to the protocol may address some of the aforementioned challenges. Introduction of backbone flexibility may significantly improve complex prediction based on unbound structures as was reported for various docking approaches^45–53^. Incorporation of additional biological constraints such as the location of the target lysine for ubiquitination, or known mutations that enhance/decrease complex formation may further restrict the conformational space.

Despite these caveats, we present an end-to-end approach to model Target/E3 ligase/PROTAC ternary complexes. Our method leverages the advantages of the highly constrained search space of this unique problem. We demonstrated how the use of various experimental results can lead to high confidence in our prediction. This work will serve as the basis for future in-silico design of new PROTACs, and may significantly reduce the time and resources that are currently required for the design of PROTACs against a new target.

## Methods

### Preparation of input files

The PDB files were downloaded from the PDB (https://www.rcsb.org/). The chains of the E3 ligase and the target protein were cleaned from any non-protein atoms. The binders were extracted manually from the PROTAC in the PDB. In cases where the PROTAC was not modeled into the structure (6BN8, 6BN9, 6BNB), we aligned another structure(s) with the same domains and binders, and copied the binders coordinates. We added hydrogens to the binders using OpenBabel (http://openbabel.org/wiki/Main_Page). The linker smiles representation was taken from the PDB. In cases it was not modeled in the PDB, it was taken from the paper where they were reported.

### Estimating linker distance constraints

We sample the length of the PROTAC in the following fashion. For each bin starting with 1Å, with increments of 1Å, we made 200 random trials to produce a PROTAC conformation (see Supp. Files main.py and ProtacLib.py). In each trial, a random orientation of the two binders is produced, while keeping the anchors in the distance set by the bin “b”. This is done by generating a random conformation of the two binders, transforming them such that the anchors are at (0,0,0), (b,0,0), followed by a random rotation of both of them. Then, using RDKit, we attempt to create a valid PROTAC conformation based on the randomized binders orientation. Once an orientation is generated, we re-measure the distance between the two anchors, to make sure it stays close to the distance set by the bin. We sum up the number of successful conformations generated in each bin, to generate a histogram. From the histogram we take the highest value, multiply it by 0.75 and set it as a threshold. Then, we take the minimum and maximum bin values that achieve that threshold: minA, maxA. Due to some cases where only a few bins were populated, we also take the average bin value of all generated conformations plus/minus 2Å as minB, maxB. The final distance constraints for the two anchors are min(minA, minB), and max(maxA, maxB).

### Constrained global protein-protein docking

Global docking is performed with PatchDock ^30^ (see Supp. Files main.py and Utils.py), using the downloadable linux version of the program. The constraints from the previous step are incorporated in the parameters file. The output is solutions of the protein-protein docking problem, clustered by a cutoff of 2Å. Up to 1,000 global docking solutions are considered for the next step.

### Local docking of selected global docking solutions

RosettaDock local docking (see Supp. Files main.py and Rosetta.py) is used to create up to 50 local docking results for each global docking solution produced by PatchDock, treating the binders as extra residues with a fixed conformation.

### Constrained PROTAC conformation generation in the context of the docking solution

We use RDKit to produce up to 100 conformations of the PROTAC, with the constraints of the two binders to fit the local docking solution (see Supp. Files main.py and Constrain_Generation.py). Therefore, only the linker’s conformation is being sampled, in regard to a fixed conformation and position of the two binders. Due to the constraints being atom-distance based, the conformation that is generated is not aligned with the position of the binders, and another alignment step is necessary. After the alignment, a threshold of 0.5Å RMSD between each head independently and its x-ray conformation is used to ensure that the conformation of the binders is really the same as their native conformation. This is necessary because RDKit generation of constrained conformations, which is based on atom-distances, can result in conformations not fully within the desired constraints. For 6SIS, 6HAX, 6HAY and 6HR2 an extra atom was added to the E3 binder, in order to uniquely define the position of the exit vector, making sure the constrained conformation attaches the linker to the right atom.

### Modeling the PROTAC within the ternary complex

We use Rosetta’s repack ^33^ protocol to choose the best PROTAC conformation from the set generated in the previous step (see Supp. Files main.py and Constrain_Generation.py). In the repack protocol, the side-chains of the residues are allowed to switch rotamers. We supplied the constrained conformations which we generated using RDKit as rotamers for the PROTAC, using the first one as the initial rotamer. Models with a final score ≥ 0 are excluded from further analysis.

### Clustering top scoring complexes

We used the DBSCAN ^36^ clustering method, and ranked the clusters by the number of models in each cluster, assuming the highly populated clusters would represent the best solutions (see Supp. Files main.py and Clustering.py). The DBSCAN method can work either with coordinates of points in n-dimensions, or with a distance-matrix of precomputed distances between each pair of points. Since we used RMSD values of the moving chain (defined always as the target protein), we precalculated all the pairwise RMSDs between the final solutions, and input it to DBSCAN as the distance-matrix. We used 4Å RMSD as the clustering threshold. We clustered the top 200 solutions based on the interface score reported by Rosetta, out of the top 1,000 scoring final solutions, created by Rosetta’s repacking protocol. We ranked the clusters according to the number of structures in each cluster. Between clusters of the same size, the ranking was based on the average score of the final models. We define a native cluster as having at least one member whose Cα RMSD from the native conformation is lower than 4Å. The reported rank is the top ranked cluster among the native clusters.

## Supporting information

Scripts for running the protocol

## Acknowledgements

We would like to thank the Rosetta Commons, and in particular Drs. Rocco Moretti and Steven Lewis for much help with algorithmic details. N.L. is the incumbent of the Alan and Laraine Fischer Career Development Chair. N.L. would like to acknowledge funding from the Israel Science Foundation (grant no. 2462/19), The Israel Cancer Research Fund, the Israeli Ministry of Science and Technology (grant no. 3-14763), the Moross Integrated Cancer Center and the Dr. Barry Sherman institute for Medicinal Chemistry. N.L. is also supported by the Helen and Martin Kimmel Center for Molecular Design, Joel and Mady Dukler Fund for Cancer Research, the Estate of Emile Mimran and Virgin JustGiving, and the George Schwartzman Fund. D.Z. would like to acknowledge the Pearlman Grant for student-initiated research in chemistry from the Faculty of Chemistry, the Weizmann Institute.

