## Supplemental Tables and Figures for "PRosettaC: Rosetta based modeling of PROTAC mediated ternary complexes"

### Supplementary Figures

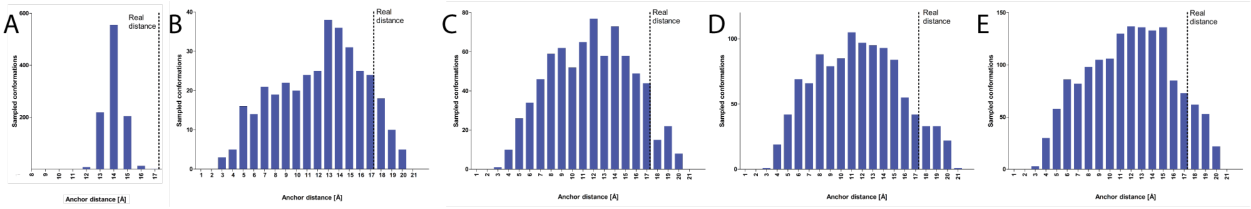


**Supplementary Figure 1. Anchor distance sampling.**
We assessed two methods to sample the distance distribution between the anchor atoms on the PROTAC target and E3 ligase binders. **A.** The most straightforward way to sample the anchor distance, is to generate random conformations of the entire PROTAC, measure the distance between the two anchor points and sum up the conformations (in bins of 1Å) to generate the distance histogram. Here we show the distribution of 100 random conformations which undersamples the actual distance for PDB: 6BOY. **B.** The alternative approach we ultimately used to estimate the distance between the anchor atoms is the following: for each value starting from 1Å and in increments of 1Å we generate random 100 pairs of binder positions with the anchor distance set to that value. Then, for each pair of binders positions, we generate a random conformation of the entire PROTAC which incorporates the 'fixed' binder positions. Due to geometrical constraints of the molecule, some of the pairs fail to generate a valid PROTAC conformation while others succeed. Once an orientation is generated, we re-measure the distance between the two anchors, to make sure it stays close to the distance set by the bin. We sum up the successfully generated PROTAC conformations for each distance bin to generate the histogram. Now the distribution covers the crystallographic distance. We select minimum and maximum distances for the protocol based on this distribution (see methods and Supp. Table 1). To assess the selection of the number of randomly generated conformations we show examples for the distribution with **C.** 200 trials **D.** 300 trials. **E.** 400 trials. The final protocol uses 200 trials to estimate the distance.

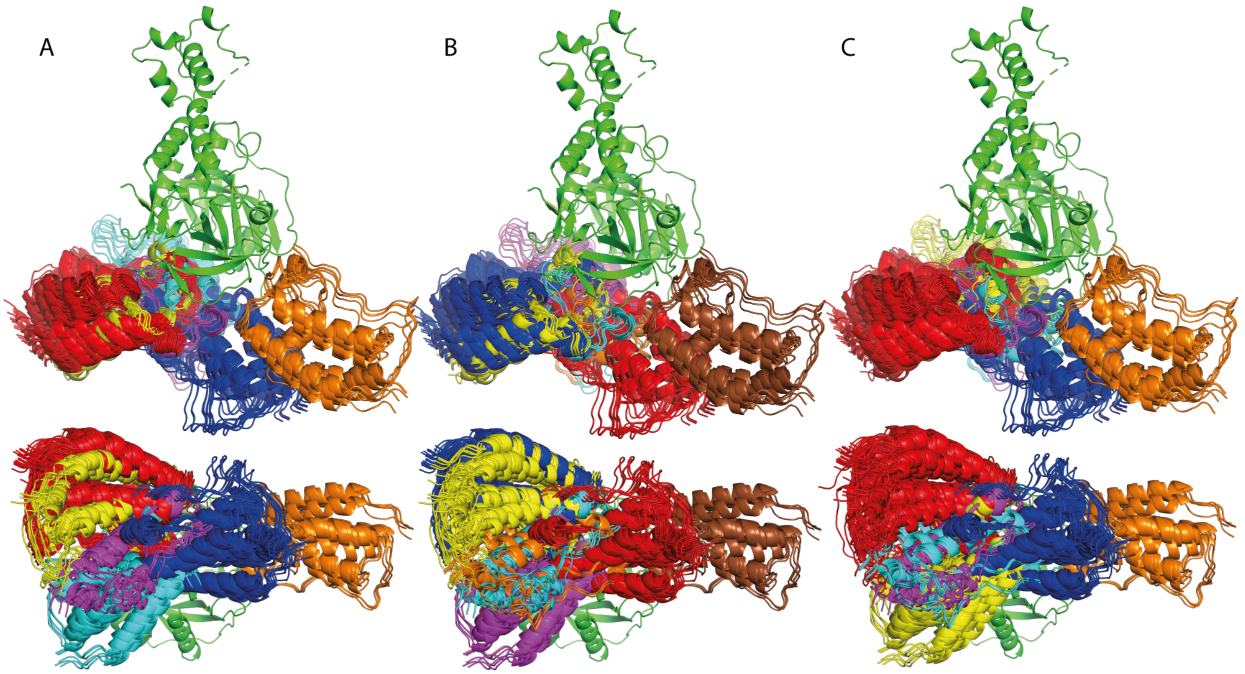


**Supplementary Figure 2. Selection of a clustering threshold.**
Different clustering thresholds for final models of PDB: 6BOY. The threshold is the neighboring distance used in the DBSCAN clustering algorithm. We chose the desired clustering threshold prior to the hyper parameters optimisation, such that the resulting clusters are coherent and represent the same binding mode. Here we only present the largest clusters, with at least five members. E3 ligase - CRBN is in green. Each cluster is presented in a different color. On the top is a side view on the protein-protein interaction, and on the bottom is a view from the bottom side of CRBN. **A.** Threshold=3Å. **B.** Threshold=4Å. **C.** Threshold=5Å. For the results reported in this manuscript we chose the threshold of 4Å.

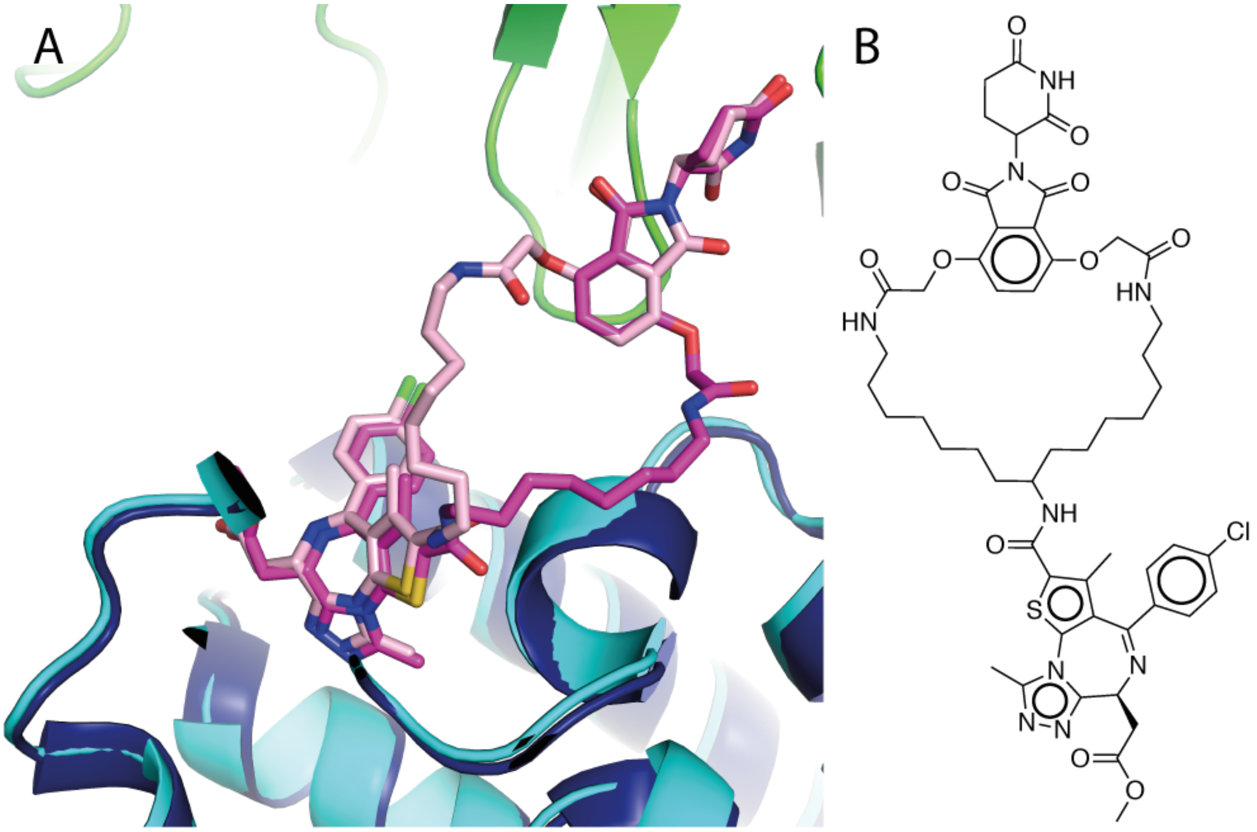


**Supplementary Figure 3. Suggested macrocyclic PROTAC based on our prediction for PDB: 6BN7.** **A.** Our top prediction (light pink) shows a possibility for the linker of 6BN7 to exit out of the opposite atom of thalidomide than the one observed in the crystal structure (magenta). Thus, we propose it may be possible to merge these linkers into a macrocyclic PROTAC, like the one developed by Tesla et al. following a similar logic. **B.** The structure of the suggested macrocyclic PROTAC.

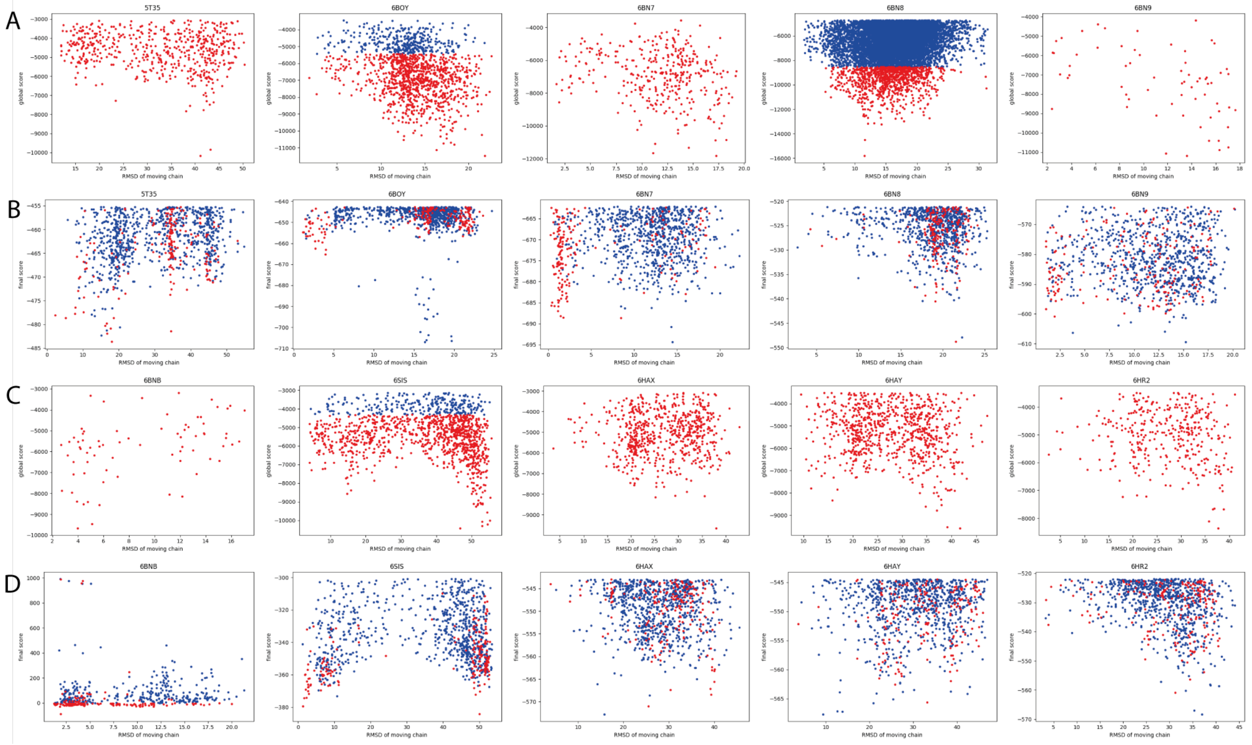


**Supplementary Figure 4. Energy landscapes of PROTAC mediated ternary-complexes.**
For each of the five cases in our 'training' set (**A & B**) and 'test' set (**C & D**) we present a plot of the energy vs. the RMSD to the crystallographic pose for models generated by global docking (**A & C**) or for final models (**B & D**).
For global docking (rows A and C), red dots represent the top 1,000 scoring solutions that were chosen, for follow-up. In case there were more than 1,000 solutions, they are colored blue. For the final models (rows B and D), red dots represent the top 200 models according to interface score that were chosen for further clustering, out of the top 1,000 scoring models by total score shown in the plot.

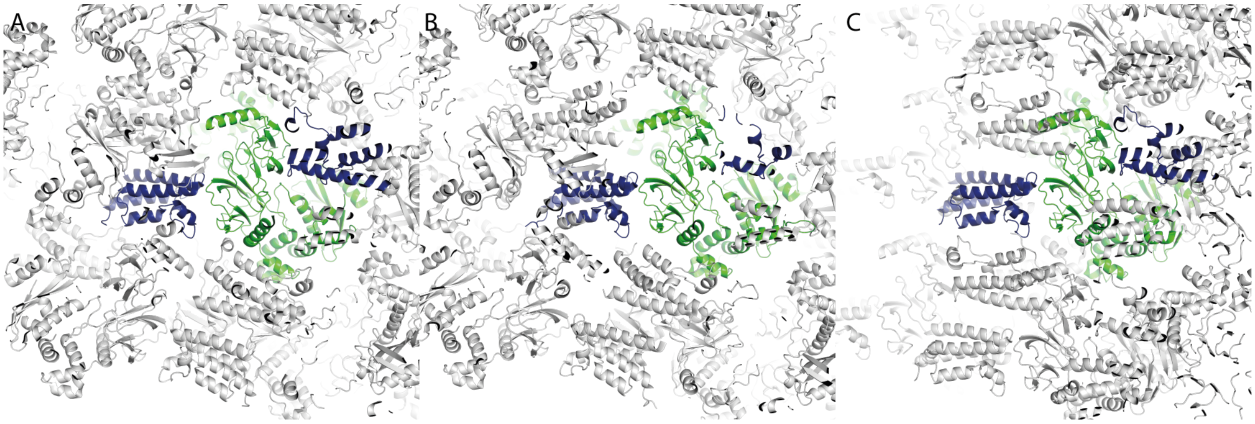


**Supplementary Figure 5. Crystal packing of VHL/SMARCA2/4 complexes.**
Crystal packing of symmetry mates (gray) of SMARCA domains (blue) in complex with VHL (green) may stabilize a specific binding conformation that is difficult for global docking to detect. **A.** PDB: 6HAX. **B.** PDB: 6HAY. **C.** PDB: 6HR2.

### Supplementary Tables

| | Train | | | | | Test | | | | |
|---|---|---|---|---|---|---|---|---|---|---|
| **Structure** | **5T35** | **6BOY** | **6BN7** | **6BN8** | **6BN9** | **6BNB** | **6SIS** | **6HAX** | **6HAY** | **6HR2** |
| **Real anchor distance (Å)** | 8.7 | 17.3 | 13.4 | 17.3 | 13.4 | 8.8 | 8.2 | 6.9 | 7.4 | 6.9 |
| **Range of anchor distance (Å)** | 6.9-13 | 8-15 | 9-15 | 10-24.3 | 7-11.7 | 4-8.2 | 5.5-11 | 4.8-8.8 | 5.4-10 | 4.7-8.7 |
| **Total Number of global docking solutions** | 555 | 1367 | 366 | 10161 | 65 | 74 | 1272 | 648 | 788 | 359 |
| **Number of global docking solutions selected for local docking** | 555 | 1000 | 366 | 1000 | 65 | 74 | 1000 | 648 | 788 | 359 |
| **Number of local docking solutions** | 17517 | 36103 | 13530 | 38243 | 2704 | 2102 | 32106 | 21143 | 26320 | 13012 |
| **Number of conformation ensembles generated** | 8065 | 27805 | 10686 | 37696 | 1872 | 527 | 4061 | 7213 | 11789 | 4778 |

**Supplementary Table 1. Statistics of distance constraints and number or models generated**
The number of conformation ensembles generated is the number of local docking solutions, for which at least one constrained conformation was generated. For the rest of local docking solutions, for which no conformations were generated after 100 trials, we assume to be incompatible with the PROTAC distance constraints and discard it.

| First round[a] | | | | | | | | | | | |
|---|---|---|---|---|---|---|---|---|---|---|---|
| PatchDock cluster size | 2 | | | | | 4 | | | | | |
| Num of local docking | 5T35 | 6BOY | 6BN7 | 6BN8 | 6BN9 | 5T35 | 6BOY | 6BN7 | 6BN8 | 6BN9 | |
| 5 | None | 3 | 3 | None | 2 | None | 3 | 4 | None | 1 | Complex |
| 10 | None | 3 | 1 | None | None | None | 37 | 1 | None | 14 | |
| 5 | None | 3 | 3 | None | 2 | None | 3 | 9 | None | 9 | Monomer |
| 10 | None | 2 | 1 | None | 3 | 18 | None | 1 | None | 1 | |

| Second round[b] | | | | | |
|---|---|---|---|---|---|
| 20 (Monomer) | 5 | 3 | 1 | None | 4 |
| 50 (Monomer) | 5 | 2 | 1 | None | 2 |

Third round[c]

| | 1000 | | | | | 2000 | | | | | 3000 | | | | |
|---|---|---|---|---|---|---|---|---|---|---|---|---|---|---|---|
| Local score | 5T35 | 6BOY | 6BN7 | 6BN8 | 6BN9 | 5T35 | 6BOY | 6BN7 | 6BN8 | 6BN9 | 5T35 | 6BOY | 6BN7 | 6BN8 | 6BN9 |
| 50 | 4 | None | 1 | None | 12 | 3 | None | 1 | None | 12 | 3 | None | 1 | None | 12 |
| 100 | 5 | 2 | 1 | None | 2 | 3 | None | 1 | None | 6 | 4 | None | 1 | None | 6 |
| 200 | 2 | 2 | 1 | None | 2 | 5 | 2 | 1 | None | 4 | 4 | 3 | 1 | None | 4 |
| 300 | 3 | 3 | 1 | None | 2 | 3 | 3 | 1 | None | 2 | 3 | 3 | 1 | None | 2 |
| 400 | 4 | 3 | 1 | None | 2 | 3 | 3 | 1 | None | 2 | 3 | 3 | 1 | None | 2 |
| 500 | 5 | 3 | 1 | None | 2 | 4 | 3 | 1 | None | 2 | 3 | 4 | 1 | None | 2 |
| 1000 | 3 | 4 | 1 | None | 1 | 8 | 3 | 2 | None | 1 | 7 | 3 | 2 | None | 1 |

| | 4000 | | | | | 5000 | | | | | 10000 | | | | |
|---|---|---|---|---|---|---|---|---|---|---|---|---|---|---|---|
| Local score | 5T35 | 6BOY | 6BN7 | 6BN8 | 6BN9 | 5T35 | 6BOY | 6BN7 | 6BN8 | 6BN9 | 5T35 | 6BOY | 6BN7 | 6BN8 | 6BN9 |
| 50 | 3 | None | 1 | None | 12 | 2 | None | 1 | None | 12 | 2 | None | 2 | None | 12 |
| 100 | 3 | None | 1 | None | 6 | 4 | None | 1 | None | 6 | 4 | None | 2 | None | 6 |
| 200 | 5 | 3 | 1 | None | 4 | 6 | 5 | 1 | None | 4 | 5 | 13 | 1 | None | 4 |
| 300 | 8 | 2 | 1 | None | 2 | 8 | 3 | 1 | None | 2 | 6 | 6 | 1 | None | 2 |
| 400 | 3 | 4 | 1 | None | 2 | 3 | 3 | 1 | None | 2 | 8 | 4 | 1 | None | 2 |
| 500 | 3 | 4 | 1 | None | 2 | 3 | 4 | 1 | None | 2 | 3 | 4 | 1 | None | 2 |
| 1000 | 6 | 4 | 1 | None | 1 | 5 | 4 | 2 | None | 1 | 6 | 4 | 2 | None | 1 |

**Supplementary Table 2. Hyper parameter optimization.**
For each condition and each of the five cases of our training set, we report the rank of the near native cluster. 'None' signifies that no near-native cluster was found.
[a] In the first round of optimization we have explored three parameters: 1. The use of the E3 ligase monomer or entire complex. 2. PatchDock clustering threshold (2Å / 4Å). 3. The number of local docking models generated for each global docking solution. (5 / 10). Other parameters (see third round below) were set to: 1,000 top scoring final models considered for clustering, 100 top models by interface score proceeding to clustering.
[b] In the second round, we fixed the clustering threshold to 2Å and the usage of the E3 monomer, and assessed if increasing the local docking sampling to 20, or 50 improved ranking, as it indeed has.
[c] In the third round we optimized the number of top scoring final models considered for clustering (1000-10,000) as well as the number of top models by interface score (50-1000) proceeding to clustering. These parameters were determined with the previously optimized parameters fixed (E3: monomer, PatchDock clustering threshold of 2Å, number of local docking decoys is 50). In all of these reported simulations, a clustering threshold of 4Å was used for the final clustering and ranking.

| | 5T35 | 6BOY | 6BN7 | 6BN8 | 6BN9 |
|---|---|---|---|---|---|
| | Drummond and Williams[1] | | | | |
| **1** | 0.0% | 0.0% | 4.3% | 0.0% | 0.0% |
| **2** | 4.4% | 0.0% | 10.0% | 2.1% | 8.1% |
| **3** | 7.9% | 0.0% | 0.0% | 0.0% | 1.0% |
| **4A** | 39.2% | 0.0% | 0.0% | 0.0% | 19.0% |
| **4B, Biased** | 39.5% | 0.0% | 8.7% | 0.0% | 13.5% |
| **4C** | 29.0% | 0.0% | 0.0% | 0.0% | 15.6% |
| **4D, Biased** | 27.0% | 0.0% | 9.1% | 0.0% | 15.5% |
| **4E, Biased** | 13.7% | 0.0% | 4.8% | 2.2% | 10.4% |
| **4F, Biased** | 12.1% | 0.0% | 4.0% | 1.9% | 11.4% |
| | PRosettaC | | | | |
| **Cluster rank** | 2/46 | 2/22 | 1/29 | - | 2/40 |
| **RMSD < 10A** | 3.5% | 17.0% | 62.0% | 0.5% | 22.0% |

**Supplementary Table 3. Comparison of PRosettaC to previously reported methods.**
Drummond and Williams[1] reported percent of models with <10Å RMSD over the E3 ligase as the moving chain for various methods. To compare PRosettaC to their methods we realigned the final 200 chosen structures (by interface score) on the target chain, and then calculated the RMSD of the E3 chain. We report here the rank of the best cluster by PRosettaC as well as the percent of models with <10Å RMSD over the E3 ligase.


1. Drummond, M. L.; Williams, C. I. In Silico Modeling of PROTAC-Mediated Ternary Complexes: Validation and Application. *J. Chem. Inf. Model.* **2019**, *59* (4), 1634–1644.

| PROTAC ID | 1 | 2 | 3 | 4 | 5 | 6 | 7 | 8 | 9 | 10 | 11 |
|---|---|---|---|---|---|---|---|---|---|---|---|
| # Structures[a] | 0 | 0 | 0 | 18 | 73 | 2527 | 7250 | 8542 | 20372 | 21107 | 20343 |

**Supplementary Table 4. Modeling a series of PROTACs against BTK.**
[a] We applied the PRosettaC protocol to BTK PROTACs **1-11** reported by Zorba et al.[2]. We report the number of final structures that were generated, before selection of top scoring structures for clustering. There is a clear distinction between PROTACs 1-4 which were inactive to PROTACs 6-11 which showed efficient degradation and formation of ternary complexes.

2. Zorba, A.; Nguyen, C.; Xu, Y.; Starr, J.; Borzilleri, K.; Smith, J.; Zhu, H.; Farley, K. A.; Ding, W.; Schiemer, J.; Feng, X.; Chang, J. S.; Uccello, D. P.; Young, J. A.; Garcia-Irrizary, C. N.; Czabaniuk, L.; Schuff, B.; Oliver, R.; Montgomery, J.; Hayward, M. M.; Coe, J.; Chen, J.; Niosi, M.; Luthra, S.; Shah, J. C.; El-Kattan, A.; Qiu, X.; West, G. M.; Noe, M. C.; Shanmugasundaram, V.; Gilbert, A. M.; Brown, M. F.; Calabrese, M. F. Delineating the Role of Cooperativity in the Design of Potent PROTACs for BTK. *Proc. Natl. Acad. Sci. U. S. A.* **2018**, *115* (31), E7285–E7292.
