## Supplementary material for "PRosettaC: Rosetta based modeling of PROTAC mediated ternary complexes": Scripts for running the protocol: Clustering.py

```
import os,sys,shutil
import ProtacLib as pl
import Rosetta as rs
import Utils
import glob
import math
from sklearn.cluster import DBSCAN
import numpy as np
sys.path.append(os.environ["COVALIB"])
from Code import *

def main(name, argv):
        if not len(argv) == 4:
                print_usage(name)
                return

        #Retrieving top resutls
        top_sc = int(argv[0])
        top_local = int(argv[1])
        RMSD = float(argv[2])
        chain = argv[3]
        os.chdir('Patchdock_Results')
        with open('score.sc', 'r') as f:
                local_lines = [line.split() for line in f][2:]
        local_lines = [line for line in local_lines if float(line[1]) < 0]
        local_lines.sort(key=lambda x: float(x[1]))
        local_lines = local_lines[:top_sc]
        top_files = [((line[-1] + '.pdb').split('_'), float(line[1])) for line in local_lines]
        for (i, j) in top_files:
                i[2] = '%04d' % (int(i[2]),)
        top_files = [('pd.' + i[1] + '_docking_' + i[2], j) for (i, j) in top_files]
        local_files = []
        for (i, j) in top_files:
                with open('local.fasc', 'r') as local:        
                        for line in local:
                                if i in line:
                                        tmp_line = line.split()
                        local_files.append((tmp_line, j))
        local_files.sort(key=lambda x: float(x[0][5]))
        local_files = [(x[-1].split('.')[1].split('_'), y) for (x, y) in local_files[:top_local]]
        local_files = [('combined_' + x[0] + '_' + str(int(x[2])) + '_0001.pdb', y) for (x, y) in local_files]
        top_folder = '../Top_RMSD' + str(RMSD) + '_' + str(top_local) + '_ISC_Top_' + str(top_sc) + '/'
        os.mkdir(top_folder)
        for f in local_files:
                shutil.copyfile(f[0], top_folder + f[0])
        #Clustering
        os.chdir(top_folder)
        shutil.copyfile('../Init.pdb', './Init.pdb')
        rank, big_clusters = apply_DBSCAN('Init.pdb', local_files, chain, RMSD)
        with open('rank.txt', 'w') as f:
                f.write(str(rank) + '\n')
        with open('big_clusters.txt', 'w') as f:
                f.write(str(big_clusters) + '\n')
        os.chdir('../')

def apply_DBSCAN(native, names, chain, threshold, cls_size_threshold=5):
	names_dict = {}
	for (x, y) in names:
		names_dict[x] = y
        with open(native, 'r') as f:
                native = [line[27:].split()[:3] for line in f if line[:4] == 'ATOM' and line[21] == chain and '  CA  ' in line]
        native = [[float(x) for x in line] for line in native]
        models = []
        for (l, y) in names:
                with open(l, 'r') as f:
                        final = [line[27:].split()[:3] for line in f if line[:4] == 'ATOM' and line[21] == chain and '  CA  ' in line]
                final = [[float(x) for x in line] for line in final]
                models.append(final)
                if not len(native) == len(final):
                        print "Not the same length"
                        sys.exit()
        num = len(models)
        rmsd = []
        for i,m1 in enumerate(models + [native]):
                line = [0]*num
                for j,m2 in enumerate(models):
                        for k in range(len(m1)):
                                for l in [0,1,2]:
                                        line[j] += (m1[k][l]-m2[k][l])**2
                        line[j] = math.sqrt(line[j]/len(m1)) 
                if i == len(models):
                        native = line
                else:
                        rmsd.append(line)
        rmsd = np.array(rmsd)
        clustering = DBSCAN(eps=threshold, metric='precomputed', min_samples=1).fit(rmsd)
        labels = clustering.labels_
        num_labels = max(labels) + 1
        native_clusters = [False]*num_labels
        cluster_size = [0]*num_labels
	cluster_sum = [0.0]*num_labels
        big_clusters = 0
        for i in range(num_labels):
                os.mkdir('tmp' + str(i + 1))
        for i,label in enumerate(labels):
                if not label == -1:
                        if native[i] <= threshold:
                                native_clusters[label] = True
                        cluster_size[label] += 1
			cluster_sum[label] += names_dict[names[i][0]]
                        shutil.copyfile('../Patchdock_Results/' + names[i][0], 'tmp' + str(label + 1) + '/' + names[i][0])
                        os.remove(names[i][0])

	cluster_avg = []
	for i in range(num_labels):
		cluster_avg.append(-1 * cluster_sum[i] / cluster_size[i])

        #Ranking the clusters, primarily by cluster size, then by average of final score
        clusters = zip(cluster_size, cluster_avg, native_clusters, ['tmp' + str(i + 1) for i in range(num_labels)])
        sorted_clusters = sorted(clusters, reverse=True)
        rank = None
        for i,c in enumerate(sorted_clusters):
                if rank == None and c[2]:
                        rank = i + 1
                os.rename(c[3], 'cluster' + str(i + 1))
                if c[0] >= cls_size_threshold:
                        big_clusters += 1
                with open('cluster' + str(i + 1) + '/avg.txt', 'w') as f:
                        f.write(str(-1 * c[1]) + '\n')
        
        return rank, big_clusters

def print_usage(name):
        print "Usage : " + name + " <top_score> <top_local> <RMSD> <chain>"
if __name__ == "__main__":
    main(sys.argv[0], sys.argv[1:])
```
