## Supplementary material for "PRosettaC: Rosetta based modeling of PROTAC mediated ternary complexes": Scripts for running the protocol: Constrain_Generation.py

```
import ProtacLib as pl
import Rosetta as rs
import Utils
import sys,os

def main(name, argv):
        if not len(argv) == 7:
                print_usage(name)
                return

        sdf_file = 'confs_' + argv[3] + '.sdf'
        v_atoms_sdf = 'v_' + argv[3] + '.sdf'
        docked_pdb = 'docked_' + argv[3] + '.pdb'
        docked_sdf = 'docked_' + argv[3] + '.sdf'
        combined = 'combined_' + argv[3] + '.pdb'
        #extracting the binders (heads)
        os.system('grep \'HETATM\' ' + argv[4] + ' > ' + docked_pdb)
        Utils.pdb2sdf(docked_pdb, docked_sdf)
        #generatign constrained conformations including 3 virtual atoms
        NBR, v_atoms_sdf = pl.GenConstConf(argv[:2], docked_sdf, argv[2], sdf_file, argv[6], v_atoms_sdf)
        #if no conformations were not able to be generated, the local docking solution is discarded
        #if they were generated, we run mol_to_params, add the generated pdb to the structure, and run relax, allowing the packer to choose the best PROTAC conformation.
        if os.path.getsize(sdf_file) == 0:
                os.remove(sdf_file)
                print "No conformations were generated"
        else:
                pdb, params = rs.mol_to_params(sdf_file, 'PTC', 'PT_' + argv[3], overwrite = False, conformers = True, nbr = NBR, v_atoms_sdf = v_atoms_sdf)
                rs.clean(argv[4], argv[5])
                os.system('cat ' + pdb + ' ' + argv[4].split('.pdb')[0] + '_' + argv[5] + '.pdb > ' + combined)
                rs.relax(combined, params, True)

def print_usage(name):
        print "Usage : <HeadA> <HeadB> <Head_Linkers> <Output_suffix> <Struct> <Chains> <Anchor_A>"
if __name__ == "__main__":
    main(sys.argv[0], sys.argv[1:])
```
