## Supplementary material for "PRosettaC: Rosetta based modeling of PROTAC mediated ternary complexes": Scripts for running the protocol: main.py

```
import os,sys,shutil
import ProtacLib as pl
import Rosetta as rs
import Utils
import glob
sys.path.append(Utils.SCRIPTS_FOL + 'PBS/')
import Cluster

def main(name, argv):
        if not len(argv) == 1:
                print_usage(name)
                return

        log = open('log.txt', 'w', 0)
        cluster = Cluster.Cluster()
        log.write('Processing inputs\n')
        with open(argv[0], 'r') as f:
                Structs = f.readline().split(':')[1].split()
                Chains = f.readline().split(':')[1].split()
                Heads = f.readline().split(':')[1].split()
                Anchors = [int(i) - 1 for i in f.readline().split(':')[1].split()]
                Linkers = f.readline().split(':')[1].split()[0]
        log.write('Cleaning structures, adding hydrogens to binders and running relax\n')
        PT_params = []
        for i in [0, 1]:
                #Adding hydrogens to the heads (binders)
                new_head = Heads[i].split('.')[0] + "_H.sdf"
                Utils.addH_sdf(Heads[i], new_head)
                Anchors[i] = pl.translate_anchors(Heads[i], new_head, Anchors[i])
                Heads[i] = new_head
                #Cleaning the structures
                rs.clean(Structs[i], Chains[i])
                Structs[i] = Structs[i].split('.')[0] + '_' + Chains[i] + '.pdb'
                #Relaxing the initial structures
                PT_pdb, PT_param = rs.mol_to_params(Heads[i], 'PT' + str(i), 'PT' + str(i))
                PT_params.append(PT_param)
                os.system('cat ' + PT_pdb + ' ' + Structs[i] + ' > Side' + str(i) + '.pdb')
                Structs[i] = 'Side' + str(i) + '.pdb'
                rs.relax(Structs[i], PT_param)
                Structs[i] = 'Side' + str(i) + '_0001.pdb'
                #Fix the atom order by adding the original ligands to the relaxed structures
                rs.clean(Structs[i], Chains[i])
                Structs[i] = Structs[i].split('.')[0] + '_' + Chains[i] + '.pdb'
                os.system('cat ' + PT_pdb + ' ' + Structs[i] + ' > Init' + str(i) + '.pdb')
                Structs[i] = 'Init' + str(i) + '.pdb'
        
        #Generate up to 200 conformations of PROTAC conformations for each anchor distance within bins
        log.write('Sampling the distance between the two anchor points\n')
        (min_value, max_value) = pl.SampleDist(Heads, Anchors, Linkers)

        #PatchDock
        log.write('Running PatchDock with the constrains\n')
        Num_Results = Utils.patchdock(Structs, [a + 1 for a in Anchors], min_value, max_value, 1000, 2.0)

        #Rosetta Local Docking
        log.write('Run Rosetta local docking on the top 1000 PatchDock results\n')
        curr_dir = os.getcwd()
        os.chdir('Patchdock_Results/')
        commands = [rs.local_docking('pd.' + str(i + 1) + '.pdb', Chains[0] + 'X', Chains[1] + 'Y', curr_dir + '/' + PT_params[0], curr_dir + '/' + PT_params[1]) for i in range(Num_Results)]
        jobs = cluster.runBatchCommands(commands, mem='8000mb')
        Cluster.Cluster.wait(jobs)

        #Generating 100 constrained conformations for the entire linker based on PatchDock results
        log.write('Generating up to 100 constrained conformations for each local docking results\n')
        docking_solutions = glob.glob('*_docking_????.pdb')
        suffix = []
        for s in docking_solutions:
                suffix.append([s, s.split('.')[1].split('_')])
                suffix[-1][1] = suffix[-1][1][0] + '_' + str(int(suffix[-1][1][2]))
        commands = ['python ' + Utils.SCRIPTS_FOL + '/Constrain_Generation.py ../' + Heads[0] + ' ../' + Heads[1] + ' ../' + Linkers + ' ' + s[1] + " " + s[0] + " " + ''.join(Chains) + ' ' + str(Anchors[0]) for s in suffix]
        jobs = cluster.runBatchCommands(commands, batch_size=6, mem='4000mb')
        Cluster.Cluster.wait(jobs)
        
        #Clustering the top 200 local docking models (according to interface RMSD), out of 1000 final scoring models
        log.write('Clustering the top results\n')
        os.system('cat ../Init0.pdb ../Init1.pdb > ../Init.pdb')
        os.chdir('../')
        os.system('python ' + Utils.SCRIPTS_FOL + '/Clustering.py 1000 200 4 ' + Chains[1])
        log.write('Done\n')
        log.close()

def print_usage(name):
        print "Usage : " + name + " <params file>\n\nFile should look like this:\nStrctures: StructA.pdb StructB.pdb\nChains: ChainsA ChainsB\nHeads: HeadA.sdf HeadB.sdf\nAnchor atoms: 6 6\nLinkers: Linkers.smi\n"
        print "Note: In the structures, head should come first\n"
if __name__ == "__main__":
    main(sys.argv[0], sys.argv[1:])
```
