## Supplementary material for "PRosettaC: Rosetta based modeling of PROTAC mediated ternary complexes": Scripts for running the protocol: chemfarm_job_batch.py

```
sendjob_text = ['#!/bin/bash\n',
'#PBS -q medium\n',
'#PBS -N un_sub_l\n',
'#PBS -l select=1:ncpus=',':mem=', '\n',
'#PBS -j eo\n',
'source /etc/pbs.conf\n',
'echo `hostname`\n',
'cd $PBS_O_WORKDIR\n']
```
