## Supplementary material for "PRosettaC: Rosetta based modeling of PROTAC mediated ternary complexes": Scripts for running the protocol: Cluster.py

```
import subprocess
import os
import sys
import PyUtils
import socket
import chemfarm_job_submission as cjob
import chemfarm_job_batch as cbatch
from subprocess import Popen, PIPE
import time

PBS_HOME = os.environ["PBS_HOME"]

class Cluster:
	def __init__(self):
		self.typ = "CHEM"
        @staticmethod
        def wait(job_ids, timeout = -1):
                jobs_done = False
                i = 0
                while not jobs_done:
                        if i == timeout:
                                break
                        time.sleep(10)
                        all_done = True
                        p = Popen([PBS_HOME + 'qstat'], stdout=PIPE, stderr=PIPE, stdin=PIPE)
                        jobs_running = [line.split()[0] for line in p.stdout.read().split('\n') if 'pbs' in line]
                        for line in job_ids:
                                if line in jobs_running:
                                        all_done = False
                        if all_done:
                                jobs_done = True
                        else:
                                print "Not done yet"
                        i += 1

	def runSingle(self, command):
		if(self.typ == "WEXAC"):
			subprocess.call(["bsub", "-u", "/dev/null", "-R", "rusage[mem=1024]", "-q", "new-all.q", "-o", "out", "-e", "err", command])
		if(self.typ == "CHEM"):
			job_file = "job_submission.sh"
			cur_job = open(job_file, 'w')
			for line in cjob.sendjob_text[:7]:
				cur_job.write(line)
			cur_job.write(cjob.sendjob_text[7] + '\'' + command + '\'')
			for line in cjob.sendjob_text[8:]:
				cur_job.write(line)
			cur_job.close()
			#subprocess.call(["/gpopt/altair/pbs/default/bin/qsub", job_file])
                        p = Popen(["/opt/pbs/bin/qsub", job_file], stdout=PIPE, stderr=PIPE, stdin=PIPE)
                        return [line.split()[0] for line in p.stdout.read().split('\n') if 'pbs' in line][0]
        def runSingleDepend(self, command, depend_jobs_file):
                with open(depend_jobs_file, 'r') as f:
                        depend_jobs = [line.split()[0] for line in f]
                if(self.typ == "WEXAC"):
                        subprocess.call(["bsub", "-u", "/dev/null", "-R", "rusage[mem=1024]", "-q", "new-all.q", "-o", "out", "-e", "err", command])
                if(self.typ == "CHEM"):
                        job_file = "job_submission.sh"
                        cur_job = open(job_file, 'w')
                        for line in cjob.sendjob_text[:1]:
                                cur_job.write(line)
                        for line in depend_jobs:
                                cur_job.write('#PBS -W depend=afterany:' + ":".join(depend_jobs))
                        for line in cjob.sendjob_text[1:7]:
                                cur_job.write(line)
                        cur_job.write(cjob.sendjob_text[7] + '\'' + command + '\'')
                        for line in cjob.sendjob_text[8:]:
                                cur_job.write(line)
                        cur_job.close()
                        subprocess.call([PBS_HOME + "qsub", job_file])

	def runSingleShell(self, command):
		job_file = "job_submission.sh"
		cur_job = open(job_file, 'w')
		for line in cjob.sendjob_text[:7]:
			cur_job.write(line)
		cur_job.write(command + '\n')
		for line in cjob.sendjob_text[9:]:
			cur_job.write(line)
		cur_job.close()
		subprocess.call([PBS_HOME + "qsub", job_file])
	def runCommands(self, commands):
		jobs = []
                for command in commands:
			jobs.append(self.runSingle(command))
                return jobs
	def runDirSingle(self, dirname, command):
		PyUtils.create_folder(dirname[:-1])
		curr = os.getcwd()
		os.chdir(dirname[:-1])
		job = self.runSingle(command)
		os.chdir(curr)
                return job
	def runDirCommands(self, dirlist, commands):
		self.checkList(dirlist, commands)
		f = open(dirlist, 'r')
		lines = f.readlines()
                jobs = []
		for dirname, command in zip(lines, commands):
			jobs.append(self.runDirSingle(dirname, command))
		f.close()
                return jobs
	def runJobs(self, dirlist, command):
		f = open(dirlist, 'r')
		lines = f.readlines()
		commands = [command] * len(lines)
		f.close()
		return self.runDirCommands(dirlist, commands)
        def runBatchJobs(self, dirlist, command, batch_size=12, mem='2000mb'):
                curr = os.getcwd()
                f = open(dirlist, 'r')
                lines = [line[:-1] for line in f.readlines()]
                f.close()
                jobs = []
                batch_list = [lines[i:i+batch_size] for i in range(0, len(lines), batch_size)]
                for i, batch in enumerate(batch_list):
                        job_file = "job_batch_" + str(i) + ".sh"
                        cur_job = open(job_file, 'w')
                        for line in cbatch.sendjob_text[:4]:
                                cur_job.write(line)
                        cur_job.write(str(len(batch)))
                        cur_job.write(cbatch.sendjob_text[4])
                        cur_job.write(mem)
                        for line in cbatch.sendjob_text[5:]:
                                cur_job.write(line)
                        for dirname in batch:
                                cur_job.write('cd ' + dirname + '\n')
                                cur_job.write(command + ' &\n')
                                cur_job.write('cd ' + curr + '\n')
                        cur_job.write('wait\n')
                        cur_job.close()
                        p = Popen(["qsub", job_file], stdout=PIPE, stderr=PIPE, stdin=PIPE)
                        jobs.append([line.split()[0] for line in p.stdout.read().split('\n') if 'pbs' in line][0])
                return jobs
        def runBatchCommands(self, commands, batch_size=12, mem='2000mb'):
                jobs = []
                batch_list = [commands[i:i+batch_size] for i in range(0, len(commands), batch_size)]
                for i, batch in enumerate(batch_list):
                        job_file = "job_batch_" + str(i) + ".sh"
                        cur_job = open(job_file, 'w')
                        for line in cbatch.sendjob_text[:4]:
                                cur_job.write(line)
                        cur_job.write(str(len(batch)))
                        cur_job.write(cbatch.sendjob_text[4])
                        cur_job.write(mem)
                        for line in cbatch.sendjob_text[5:]:
                                cur_job.write(line)
                        for command in batch:
                                cur_job.write(command + ' &\n')
                        cur_job.write('wait\n')
                        cur_job.close()
                        p = Popen(["qsub", job_file], stdout=PIPE, stderr=PIPE, stdin=PIPE)
                        jobs.append([line.split()[0] for line in p.stdout.read().split('\n') if 'pbs' in line][0])
                return jobs
        def runJobsArgs(self, dirlist, command, arglist):
		self.checkList(dirlist, arglist)
		commands = []
		for arg in arglist:
			commands += [command + ' ' + str(arg)]
		self.runDirCommands(dirlist, commands)
	
	def runCommandsArgs(self, command, arglist):
		commands = []
		for arg in arglist:
			commands.append(command + ' ' + str(arg))
		return self.runCommands(commands)
        def runJobsName(self, dirlist, command):
                with open(dirlist) as f:
                        lines = f.read().splitlines()
                self.runJobsArgs(dirlist, command, lines)

	#Inner functions
	def getServer(self):
		server_name = socket.gethostname()
		if 'chemfarm' in server_name:
			self.typ = "CHEM"
		elif 'wexac' in server_name:
			self.typ = "WEXAC"
		else:
			print "This is not a cluster supporting server"
			sys.exit()
	def checkList(self, dirlist, l):
		f = open(dirlist, 'r')
		if not len(f.readlines()) == len(l):
			print 'length of list does not match dirlist length'
			sys.exit()
		f.close()
```
