## Supplementary material for "PRosettaC: Rosetta based modeling of PROTAC mediated ternary complexes": Scripts for running the protocol: PyUtils.py

```
import os
def create_folder(name):
	if(os.path.exists(name) == False):
		os.mkdir(name)
def create_softlink(old, new):
        if(os.path.exists(new) == False):
                os.symlink(old, new)
def getNcolumn(file_name, n):
	Ncolumn = []
	for line in open(file_name):
		columns = line.split()
		if(len(columns) >= n + 1):
    			Ncolumn.append(columns[n])
    	return Ncolumn
def initPythonVs():
	os.system("module load python")
```
