## Supplementary material for "PRosettaC: Rosetta based modeling of PROTAC mediated ternary complexes": Scripts for running the protocol: ProtacLib.py

```
import sys
from rdkit import Chem
from rdkit.Chem import AllChem
from rdkit.Chem import rdFMCS
from rdkit.Chem import rdMolAlign
from rdkit.Chem import rdMolTransforms
from rdkit.Geometry.rdGeometry import Point3D
from rdkit.Chem.rdchem import Atom
import numpy as np
import math
import glob
import random
import copy

def translate_anchors(old_sdf, new_sdf, old_anchor):
    OldSdf = Chem.SDMolSupplier(old_sdf)[0]
    NewSdf = Chem.SDMolSupplier(new_sdf)[0]
    NewMatch = NewSdf.GetSubstructMatch(OldSdf)
    return NewMatch[old_anchor]

def rmsd(query, ref, q_match, r_match):
    rmsd = 0
    for i in range(len(q_match)):
        rmsd += (query.GetConformer().GetAtomPosition(q_match[i])-ref.GetConformer().GetAtomPosition(r_match[i])).LengthSq()
    rmsd = np.sqrt(rmsd/len(q_match))
    return rmsd

def heads_rmsd(query, headA, headB, headA_sub, headB_sub, match_A, match_B):
    rmsd = 0
    for i in range(len(match_A)):
        rmsd += (query.GetConformer().GetAtomPosition(match_A[i])-headA.GetConformer().GetAtomPosition(headA_sub[i])).LengthSq()
    for i in range(len(match_B)):
        rmsd += (query.GetConformer().GetAtomPosition(match_B[i])-headB.GetConformer().GetAtomPosition(headB_sub[i])).LengthSq()
    rmsd = np.sqrt(rmsd/(len(match_A) + len(match_B)))
    return rmsd

def MCS_AtomMap(query, ref):
    mcs = rdFMCS.FindMCS([query, ref], ringMatchesRingOnly=True, completeRingsOnly=True)
    mcs_patt = Chem.MolFromSmarts(mcs.smartsString)
    refMatch = ref.GetSubstructMatch(mcs_patt)
    queryMatch = query.GetSubstructMatch(mcs_patt)
    amap = []
    for i in range(len(refMatch)):
        amap.append((queryMatch[i], refMatch[i]))
    return amap

def SetCoordsForMatch(query, ref, ref_match):
    for i, pos in enumerate(ref_match):
        ref_pos = ref.GetConformer().GetAtomPosition(i)
        query.GetConformer().SetAtomPosition(pos, ref_pos)

def translateMol(mol, pointA, pointB):
    for i in range(mol.GetConformer().GetNumAtoms()):
        mol.GetConformer().SetAtomPosition(i, mol.GetConformer().GetAtomPosition(i) + pointA - pointB)

def x_rotation(vector,theta):
    """Rotates 3-D vector around x-axis"""
    R = np.array([[1,0,0],[0,np.cos(theta),-np.sin(theta)],[0, np.sin(theta), np.cos(theta)]])
    output = np.dot(R, [vector.x, vector.y, vector.z])
    return Point3D(output[0], output[1], output[2])

def y_rotation(vector,theta):
    """Rotates 3-D vector around y-axis"""
    R = np.array([[np.cos(theta),0,np.sin(theta)],[0,1,0],[-np.sin(theta), 0, np.cos(theta)]])
    output = np.dot(R, [vector.x, vector.y, vector.z])
    return Point3D(output[0], output[1], output[2])

def z_rotation(vector,theta):
    """Rotates 3-D vector around z-axis"""
    R = np.array([[np.cos(theta), -np.sin(theta),0],[np.sin(theta), np.cos(theta),0],[0,0,1]])
    output = np.dot(R, [vector.x, vector.y, vector.z])
    return Point3D(output[0], output[1], output[2])

#rotate a molecule around the origin
def rotateMol(mol, x, y, z):
    for i in range(mol.GetConformer().GetNumAtoms()):
        atom = mol.GetConformer().GetAtomPosition(i)
        atom = x_rotation(atom, x)
        atom = y_rotation(atom, y)
        atom = z_rotation(atom, z)
        mol.GetConformer().SetAtomPosition(i, atom)

def randomRotateMol(mol):
    x = random.random()*np.pi*2
    y = random.random()*np.pi*2
    z = random.random()*np.pi*2
    rotateMol(mol, x, y, z)

#sample an array of distances between the anchor points
def SampleDist(Heads, Anchors, Linkers, n = 200, output_hist="initial_distances.hist", hist_threshold = 0.75, min_margin = 2, homo_protac = False):
    writer = Chem.SDWriter("random_sampling.sdf")
    random.seed(0)
    [HeadA_sdf, HeadB_sdf] = Heads
    #linkers                                                                                                                                  
    with open(Linkers, 'r') as f:
        linkers = [Chem.MolFromSmiles(line.split()[0]) for line in f]
    #loading the heads sdf files                                                                                                              
    HeadA = Chem.SDMolSupplier(HeadA_sdf)[0]
    HeadB = Chem.SDMolSupplier(HeadB_sdf)[0]
    origin = Point3D(0,0,0)
    anchor_a = HeadA.GetConformer().GetAtomPosition(Anchors[0])
    translateMol(HeadA, origin, anchor_a)
    anchor_b = HeadB.GetConformer().GetAtomPosition(Anchors[1])
    translateMol(HeadB, origin, anchor_b)
    for linker in linkers:
        #homo protacs are protacs with the same binder twice, causing self degradation of an E3 ligase
        if homo_protac:
            head_A = linker.GetSubstructMatches(HeadA)[0]
            head_B = linker.GetSubstructMatches(HeadB)[1]
        else:
            head_A_list = linker.GetSubstructMatches(HeadA, uniquify=False)
            head_B_list = linker.GetSubstructMatches(HeadB, uniquify=False)
        histogram = {}
        seed  = 0
        b = 1
        while True:
            b_counter = 0
            for i in range(n):
                head_A = random.choice(head_A_list)
                head_B = random.choice(head_B_list)
                seed += 1
                NewA = copy.deepcopy(HeadA)
                NewB = copy.deepcopy(HeadB)
                randomRotateMol(NewA) 
                randomRotateMol(NewB)
                translateMol(NewB, Point3D(b, 0, 0), origin)
                #the constraints for the conformation generation using the two randomized heads                                               
                cmap = {head_A[i]:NewA.GetConformer().GetAtomPosition(i) for i in range(len(head_A))}
                cmap.update({head_B[i]:NewB.GetConformer().GetAtomPosition(i) for i in range(len(head_B))})
                #only half of the atoms are required to make the constrained embedding
                #this is done because using all the atoms sometimes makes it impossible
                #to find solutions, the half is chosen randomly for each generation
                cmap_tag = random.sample(cmap, len(cmap)/2)
                cmap_tag = {ctag:cmap[ctag] for ctag in cmap_tag}
                if AllChem.EmbedMolecule(linker, coordMap=cmap_tag, randomSeed=seed, useBasicKnowledge=True, maxAttempts=1) == -1:
                    continue
                if int(round(rdMolTransforms.GetBondLength(linker.GetConformer(), head_A[Anchors[0]], head_B[Anchors[1]]))) == b:
                    writer.write(linker)
                    b_counter += 1
            histogram[b] = b_counter
            if b >= 10 and b_counter == 0:
                break
            b += 1
        with open(output_hist, 'w') as f:
            for h in histogram:
                f.write(str(h) + "\t" + str(histogram[h]) + '\n')
        max_value = max([histogram[i] for i in histogram])
        sum_mul = 0
        sum_his = 0
        for i in histogram:
            sum_mul += i * histogram[i]
            sum_his += histogram[i]
        avg_index = 1.0 * sum_mul / sum_his
        threshold = max_value * hist_threshold
        high_values = [i for i in histogram if histogram[i] >= threshold]
        return(min(min(high_values), avg_index - min_margin), max(max(high_values), avg_index + min_margin))

#generate n random conformations for each smile in linkers_file
#this is an old function and is not used in the final pipeline
def GenRandConf(Heads, Anchors, Linkers, n=1000, output_hist="initial_distances.hist", output_sdf="random_sampling.sdf"):
    writer = Chem.SDWriter(output_sdf)
    [HeadA_sdf, HeadB_sdf] = Heads
    #linkers
    with open(Linkers, 'r') as f:
        linkers = [Chem.MolFromSmiles(line.split()[0]) for line in f]
    #loading the heads sdf files
    HeadA = Chem.SDMolSupplier(HeadA_sdf)[0]
    HeadB = Chem.SDMolSupplier(HeadB_sdf)[0]
    #anchor distances
    X1Y1_dist = []
    for linker in linkers:
        Chem.AddHs(linker)
        amapA = MCS_AtomMap(HeadA, linker)
        amapB = MCS_AtomMap(HeadB, linker)
        anchors = [[item[1] for item in amapA if item[0] == Anchors[0]][0], [item[1] for item in amapB if item[0] == Anchors[1]][0]]
        i = 0
        seed = 0
        while i < n:
            seed += 1
            if Chem.rdDistGeom.EmbedMolecule(linker, randomSeed=seed, useBasicKnowledge=True, maxAttempts=10) == -1:
                continue
            X1Y1_dist.append(rdMolTransforms.GetBondLength(linker.GetConformer(), anchors[0], anchors[1]))
            writer.write(linker)
            i += 1
    
    hist = np.histogram(np.array(X1Y1_dist), range=(math.floor(min(X1Y1_dist)), math.ceil(max(X1Y1_dist))), bins=2*int(math.ceil(max(X1Y1_dist)-math.floor(min(X1Y1_dist)))))
    with open(output_hist, 'w') as f:
        f.write("range is: " + str([min(X1Y1_dist), max(X1Y1_dist)]) + '\n')
        for i in range(len(hist[0])):
            f.write(str(hist[0][i]) + '\t' + str(hist[1][i]) + '\n')
    return (min(X1Y1_dist), max(X1Y1_dist))

#generate constrained conformations for each of the local docking solutions
def GenConstConf(Heads, Docked_Heads, Head_Linkers, output_sdf, Anchor_A, v_atoms_sdf, n = 100, homo_protac = False):
    writer = Chem.SDWriter(output_sdf)
    with open(Head_Linkers, 'r') as f:
        head_linkers = [Chem.MolFromSmiles(line.split()[0]) for line in f]
    #loading the heads sdf files
    HeadA = Chem.SDMolSupplier(Heads[0])[0]
    HeadB = Chem.SDMolSupplier(Heads[1])[0]
    docked_heads = Chem.SDMolSupplier(Docked_Heads)[0]
    #virtual atoms around the center of mass for the neighbor atom alignment
    num_atoms = docked_heads.GetConformer().GetNumAtoms()
    x = []
    y = []
    z = []
    for i in range(num_atoms):
        x.append(docked_heads.GetConformer().GetAtomPosition(i).x)
        y.append(docked_heads.GetConformer().GetAtomPosition(i).y)
        z.append(docked_heads.GetConformer().GetAtomPosition(i).z)
    v1 = Point3D(sum(x)/num_atoms, sum(y)/num_atoms, sum(z)/num_atoms)
    v2 = Point3D(sum(x)/num_atoms + 1, sum(y)/num_atoms, sum(z)/num_atoms)
    v3 = Point3D(sum(x)/num_atoms, sum(y)/num_atoms + 1, sum(z)/num_atoms)
    virtual_atoms = Chem.MolFromSmarts('[#23][#23][#23]')
    Chem.rdDistGeom.EmbedMolecule(virtual_atoms)
    virtual_atoms.GetConformer().SetAtomPosition(1, v1)
    virtual_atoms.GetConformer().SetAtomPosition(0, v2)
    virtual_atoms.GetConformer().SetAtomPosition(2, v3)
    v_writer = Chem.SDWriter(v_atoms_sdf)
    v_writer.write(virtual_atoms)

    #homo protacs are protacs with the same binder twice, causing self degradation of an E3 ligase
    if homo_protac:
        docked_A = docked_heads.GetSubstructMatches(HeadA)[0]
        docked_B = docked_heads.GetSubstructMatches(HeadB)[1]
    else:
        docked_A = docked_heads.GetSubstructMatch(HeadA)
        docked_B = docked_heads.GetSubstructMatch(HeadB)
    for head_linker in head_linkers:
        Chem.AddHs(head_linker)
        if homo_protac:
            head_A = head_linker.GetSubstructMatches(HeadA)[0]
            head_B = head_linker.GetSubstructMatches(HeadB)[1]
        else:
            head_A_list = head_linker.GetSubstructMatches(HeadA, uniquify=False)
            head_B_list = head_linker.GetSubstructMatches(HeadB, uniquify=False)

        i = 0
        seed = 0
        while i < n:
            if seed > 10 * n:
                break
            seed += 1

            random.seed(seed)

            head_A = random.choice(head_A_list)
            head_B = random.choice(head_B_list)

            #amap for final alignment                                                                                                         
            amap = []
            for j in range(len(docked_A)):
                amap.append((head_A[j], docked_A[j]))
            for j in range(len(docked_B)):
                amap.append((head_B[j], docked_B[j]))

            #the constraints for the conformation generation using the two docked heads                                                       
            cmap = {head_A[j]:docked_heads.GetConformer().GetAtomPosition(docked_A[j]) for j in range(len(docked_A))}
            cmap.update({head_B[j]:docked_heads.GetConformer().GetAtomPosition(docked_B[j]) for j in range(len(docked_B))})

            #only half of the atoms are required to make the constrained embedding
            #this is done because using all the atoms sometimes makes it impossible
            #to find solutions, the half is chosen randomly for each generation
            cmap_tag = random.sample(cmap, len(cmap)/2)
            cmap_tag = {ctag:cmap[ctag] for ctag in cmap_tag}

            if AllChem.EmbedMolecule(head_linker, coordMap=cmap_tag, randomSeed=seed, useBasicKnowledge=True, maxAttempts=10) == -1:
                continue
            
            #final alignment to bring the new conformation to the position of the pose's heads
            #this is needed because the constrained embedding only applies
            #to distances and not to atoms position
            rdMolAlign.AlignMol(head_linker, docked_heads, atomMap=amap)
            #make sure the alignment is good enough for both heads (also to ensure the save isomer
            #for ambiguous rings
            if rmsd(head_linker, docked_heads, head_A, docked_A) < 0.5 and rmsd(head_linker, docked_heads, head_B, docked_B) < 0.5:
                writer.write(head_linker)
                i += 1
    return head_A[int(Anchor_A)], v_atoms_sdf

#printing head RMSD, calculated between a modeled PROTAC, and the .sdf files of the two binders (headA, headB)
def print_rmsd(headA, headB, Head_Linkers):
    #loading the heads sdf files                                                                                      
    HeadA = Chem.SDMolSupplier(headA)[0]
    HeadB = Chem.SDMolSupplier(headB)[0]
    head_linkers = Chem.SDMolSupplier(Head_Linkers)
    for head_linker in head_linkers:
        mcsA = rdFMCS.FindMCS([HeadA, head_linker])
        mcsB = rdFMCS.FindMCS([HeadB, head_linker])
        pattA = Chem.MolFromSmarts(mcsA.smartsString)
        pattB = Chem.MolFromSmarts(mcsB.smartsString)
        HeadA_sub = HeadA.GetSubstructMatches(pattA, uniquify=False)
        HeadB_sub = HeadB.GetSubstructMatch(pattB)
        head_A = head_linker.GetSubstructMatches(pattA, uniquify=False)
        head_B = head_linker.GetSubstructMatch(pattB)
        for H in HeadA_sub:
            for h in head_A:
                print heads_rmsd(head_linker, HeadA, HeadB, H, HeadB_sub, h, head_B)
```
