## Supplementary material for "PRosettaC: Rosetta based modeling of PROTAC mediated ternary complexes": Scripts for running the protocol: README

Installation requirements for PRosettaC:

1. Install RDKit and NumPy python packages.

2. Download and Install OpenBabel (http://openbabel.org/wiki/Main_Page), PatchDock (https://bioinfo3d.cs.tau.ac.il/PatchDock/) and Rosetta (https://www.rosettacommons.org/).

3. Our scripts rely on the PBS job scheduler (https://en.wikipedia.org/wiki/Portable_Batch_System).

4. Set the following environment parameters: export PATCHDOCK=”/Path_to_PatchDock/”, export OB=”/Path_to_PatchDock/”, export SCRIPTS_FOL=”/Path_to_the_folder_containing_the_scripts/”, export ROSETTA_FOL=”/Path_to_Rosetta/”, PBS_HOME=”/Path_to_PBS_bin_folder/”.

All of the scripts required for running the protocol are supplied as supplementary files. 

Usage: python main.py Protac_params.txt

Protac_param.txt will include the input files and should look like this:

Structures: StructA.pdb StructB.pdb
Chains: A B
Heads: HeadA.sdf HeadB.sdf
Anchor atoms: 11 23
Linkers: Linkers.smi

StructA.pdb is the structure of the E3 ligase (e.g. CRBN)
StructB.pdb is the structure of the target protein (e.g. BRD4)
A and B are the chain IDs for the E3 ligase and target. If your structures have more than one chain, write them like this: Chains: AC B. There should not be identical chain IDs for the E3 ligase and the target.
HeadA.sdf is the E3 ligase binder (e.g. thalidomide)
HeadB.sdf is the target binder (e.g. JQ1)
Both HeadA and HeadB should be positioned in their appropriate binding modes within StructA and StructB.
Anchor atoms is the number of the anchor atoms within HeadA and HeadB. An easy way to find the number of a desired atom is to open HeadA.sdf in pymol, press the L button in the toolbar, then choose "atom identifiers", and "ID". Just make sure that the anchor should be >= 1, so in case the numbering in pymol starts from 0, add one to the number of the chosen atom. The anchor atoms must be uniquely defined in the smiles representation of the molecule.
Linkers.smi is a file with a single line which is the smiles representation of the full PROTAC.

Authors:
Daniel Zaidman, Nir London
