## Supplementary material for "PRosettaC: Rosetta based modeling of PROTAC mediated ternary complexes": Scripts for running the protocol: Rosetta.py

```
import os,sys
import Utils

FOLDER = Utils.SCRIPTS_FOL + "Rosetta/"
SCRIPTS = Utils.ROSETTA_FOL + "/main/source/bin/rosetta_scripts.default.linuxgccrelease"
CLEAN = Utils.ROSETTA_FOL + "/tools/protein_tools/scripts/clean_pdb.py"

#get only the ATOM entries for specific chains
def clean(struct, chains):
    os.system(CLEAN + ' ' + struct + ' ' + chains)
    for chain in chains:
        os.remove(struct.split('.pdb')[0] + '_' + chain + '.fasta')

#replace the old structure with the clean version
def clean_replace(struct, chains):
    os.system(CLEAN + ' ' + struct + ' ' + chains)
    for chain in chains:
        os.remove(struct.split('.pdb')[0] + '_' + chain + '.fasta')
    os.rename(struct.split('.pdb')[0] + '_' + chains + '.pdb', struct)

#run the mol_to_param Rosetta script
def mol_to_params(sdf_file, name, pdb, overwrite = True, conformers = False, nbr = -1, v_atoms_sdf = ''):
    Utils.addH_sdf(sdf_file)
    if not v_atoms_sdf == '':
        nbr = Utils.add_virtual_atoms(sdf_file, v_atoms_sdf, sdf_file)
    line = "/home/danielza/Work/Software/Rosetta/main/source/scripts/python/public/molfile_to_params.py " + sdf_file + " -n " + name + " -p " + pdb
    if overwrite:
        line += " --clobber"
    if conformers:
        line += " --conformers-in-one-file"
    if not nbr == -1:
        line += " --nbr_atom=" + str(nbr)
    os.system(line)
    if conformers:
        return pdb + '.pdb', pdb + '.params'
    return pdb + '_0001.pdb', pdb + '.params'

#run a relax protocol
def relax(struct, sdf_params, interface = False, n = 1):
    if interface:
        os.system(SCRIPTS + " -s " + struct + " -parser:protocol " + FOLDER + "int_relax.xml @" + FOLDER + "relax.flags -extra_res_fa " + sdf_params + " -overwrite")
    elif n == 1:
        os.system(SCRIPTS + " -s " + struct + " -parser:protocol " + FOLDER + "relax.xml @" + FOLDER + "relax.flags -extra_res_fa " + sdf_params + " -overwrite")
    else:
        os.system(SCRIPTS + " -s " + struct + " -parser:protocol " + FOLDER + "flex_relax.xml @" + FOLDER + "relax.flags -extra_res_fa " + sdf_params + " -overwrite -nstruct " + str(n))

#run a local docking protocol
def local_docking(struct, chainsA, chainsB, ptA_params, ptB_params):
    return SCRIPTS + " -s " + struct + " -parser:protocol " + FOLDER + "docking.xml @" + FOLDER + "docking.flags @" + FOLDER + "relax.flags -extra_res_fa " + ptA_params + " -extra_res_fa " + ptB_params + " -partners " + chainsA + "_" + chainsB + " -scorefile local.fasc -overwrite"
```
