## Supplementary material for "PRosettaC: Rosetta based modeling of PROTAC mediated ternary complexes": Scripts for running the protocol: Utils.py

```
import os,sys
import glob
import shutil
PatchDock = os.environ["PATCHDOCK"]
OB = os.environ["OB"]
SCRIPTS_FOL = os.environ["SCRIPTS_FOL"]
ROSETTA_FOL = os.environ["ROSETTA_FOL"]

#adding hydrogens to an sdf file is done by converting it to pdb, and then back to sdf using openbabel
def addH_sdf(sdf_file, new_sdf = None):
    tmp_pdb = sdf_file.split('.')[0] + '.pdb'
    os.system(OB + '/babel ' + sdf_file + ' ' + tmp_pdb + ' -h')
    if new_sdf == None:
        os.system(OB + '/babel ' + tmp_pdb + ' ' + sdf_file)
    else:
        os.system(OB + '/babel ' + tmp_pdb + ' ' + new_sdf)
    os.remove(tmp_pdb)

#openbabel conversion of pdb to sdf
def pdb2sdf(pdb_file, sdf_file):
    os.system(OB + '/babel ' + pdb_file + ' ' + sdf_file)

#merge a virtual atom sdf file into an input sdf file
def add_virtual_atoms(input_sdf, v_atoms_sdf, output_sdf):
    with open(input_sdf, 'r') as f:
        confs = f.read().split('$$$$')
    confs[0] = '\n' + confs[0]
    confs = [[l + '\n' for l in line.split('\n') + ['$$$$']][1:] for line in confs][:-1]
    with open(v_atoms_sdf, 'r') as f:
        v_atoms = f.readlines()[4:7]
    header = confs[0][3]
    num_atoms = int(header[:3])
    num_bonds = int(header[3:6])
    new_header = '%03d' % (num_atoms + 3) + '%03d' % (num_bonds + 3) + header[6:]
    v_index = ['%03d' % (num_atoms + 1), '%03d' % (num_atoms + 2), '%03d' % (num_atoms + 3)]
    v_bonds = [v_index[0] + v_index[1] + '  1  0\n', v_index[1] + v_index[2] + '  1  0\n', '  1' + v_index[0] + '  1  0\n']
    for i, conf in enumerate(confs):
        confs[i] = conf[:3] + [new_header] + conf[4:4 + num_atoms] + v_atoms + conf[4 + num_atoms:4 + num_atoms + num_bonds] + v_bonds + conf[4 + num_atoms + num_bonds:-4] + ['        0    0    0    0    0    0    0    0  ' + '%03d' % (num_atoms + 3) + '    0  ' + '%03d' % (num_atoms + 3) + '    0'] + conf[-3:]
    with open(output_sdf, 'w') as f:
        for conf in confs:
            for c in conf:
                f.write(c)
    return num_atoms + 2

#run PatchDock
def patchdock(structs, anchors, min_dist, max_dist, num_results=1000, threshold=4.0):
    [structA, structB] = structs
    [anchorA, anchorB] = anchors
    with open('Patchdock_cst', 'w') as f:
        f.write(' '.join([str(s) for s in [anchorA, anchorB, min_dist, max_dist]]) + '\n')
    os.system(PatchDock + '/buildParams.pl ' + structA + ' ' + structB + ' ; mv params.txt Patchdock_params.txt')
    os.system('sed -i \'s/#distanceConstraintsFile file_name/distanceConstraintsFile Patchdock_cst/g\' Patchdock_params.txt')
    os.system('sed -i \'s/clusterParams 0.1 4 2.0 4.0/clusterParams 0.1 4 2.0 ' + str(threshold) + '/g\' Patchdock_params.txt')
    os.system(PatchDock + '/patch_dock.Linux Patchdock_params.txt Patchdock_output')
    os.system(PatchDock + '/transOutput.pl Patchdock_output 1 ' + str(num_results))
    os.mkdir('Patchdock_Results')
    Num_Results = max([int(line.split('.')[1]) for line in glob.glob('Patchdock_output.*')])
    for i in range(Num_Results):
        os.system('mv Patchdock_output.' + str(i + 1) + '.pdb Patchdock_Results/pd.' + str(i + 1) + '.pdb')
        os.system('sed -i \'s/PT1 X/PT1 Y/g\' Patchdock_Results/pd.' + str(i + 1) + '.pdb')
    return Num_Results
```
